# Whole-Genome Association Analyses of Sleep-disordered Breathing Phenotypes in the NHLBI TOPMed Program

**DOI:** 10.1101/652966

**Authors:** Brian E. Cade, Jiwon Lee, Tamar Sofer, Heming Wang, Man Zhang, Han Chen, Sina A. Gharib, Daniel J. Gottlieb, Xiuqing Guo, Jacqueline M. Lane, Jingjing Liang, Xihong Lin, Hao Mei, Sanjay R. Patel, Shaun M. Purcell, Richa Saxena, Neomi A. Shah, Daniel S. Evans, Craig L. Hanis, David R. Hillman, Sutapa Mukherjee, Lyle J. Palmer, Katie L. Stone, Gregory J. Tranah, NHLBI Trans-Omics for Precision Medicine (TOPMed) Consortium, Gonçalo R. Abecasis, Eric A. Boerwinkle, Adolfo Correa, L. Adrienne Cupples, Robert C. Kaplan, Deborah A. Nickerson, Kari E. North, Bruce M. Psaty, Jerome I. Rotter, Stephen S. Rich, Russell P. Tracy, Ramachandran S. Vasan, James G. Wilson, Xiaofeng Zhu, Susan Redline, TOPMed Sleep Working Group

## Abstract

Sleep-disordered breathing (SDB) is a common disorder associated with significant morbidity. Through the NHLBI Trans-Omics for Precision Medicine (TOPMed) program we report the first whole-genome sequence analysis of SDB. We identified 4 rare gene-based associations with SDB traits in 7,988 individuals of diverse ancestry and 4 replicated common variant associations with inclusion of additional samples (n=13,257). We identified a multi-ethnic set-based rare-variant association (p = 3.48 × 10^−8^) on chromosome X with *ARMCX3*. Transcription factor binding site enrichment identified associations with genes implicated with respiratory and craniofacial traits. Results highlighted associations in genes that modulate lung development, inflammation, respiratory rhythmogenesis and *HIF1A*-mediated hypoxic response.

## Introduction

Sleep-disordered breathing (SDB) is a prevalent disorder associated with increased mortality and morbidity [^1–4^]. The most common type of SDB is obstructive sleep apnea (OSA), characterized by repeated airway collapse leading to intermittent hypoxemia and sleep disruption, that is increaed in prevalence with older age and male sex [^5^]. The disease appears to be multifactorial, reflecting variable contributions of abnormalities in ventilatory control, craniofacial anatomy, and adiposity [^5–11^]. Due to an incomplete understanding of its pathophysiology, standard OSA treatment only addresses the downstream manifestations of airway collapse through nightly use of pressurized air to the nasopharynx, a therapy that often is poorly tolerated. Therefore, there is a critical need to identify molecular pathways that could provide specific therapeutic targets. The need for overnight studies to phenotype SDB traits has limited the available sample size for genetic analyses, and only several common-frequency genome-wide analysis studies have been reported [^11–15^]. Increased statistical power may increase the genetic resolution of regions that may not be adequately tagged by current genotyping arrays due to population differences and/or reduced linkage disequilibrium with biologically relevant regions [^16^].

The Trans-Omics for Precision Medicine (TOPMed) program is an NIH National Heart, Lung, and Blood Institute program designed to improve the understanding of the biological processes that contribute to heart, lung, blood, and sleep disorders [^17^]. TOPMed has generated whole-genome sequencing (WGS) data on over 100,000 individuals from multiple cohorts at >30× depth, including seven studies with objective assessment of SDB. A variant imputation server using TOPMed data also allows for high-quality imputation of non-sequenced genotype chip data [^18^]. A complementary initiative sponsored by the Centers for Common Disease Genomics (CCDG) of the NIH National Human Genome Research Institute has generated sequencing data from additional individuals in two TOPMed cohorts (https://www.genome.gov/27563570). These initiatives provide the ability to examine the genetics of SDB at unprecedented detail in African-Americans (AA), Asian-Americans (AsA), European-Americans/Australians (EA), and Hispanic/Latino-Americans (HA).

In this first WGS analysis of SDB, we examine the apnea-hypopnea index (AHI), the standard clinic metric of SDB, and four complementary measurements of overnight hypoxemia: average and minimum oxyhemoglobin saturation (SpO_2_) during sleep and the percent of the sleep recording with SpO_2_ < 90% (Per90); and the average desaturation per hypopnea event. These indices were chosen because of clinical relevance, high heritability, or prior significant GWAS findings [^7–9,11,14^]. We examined 7,988 individuals with objectively measured SDB and WGS data in conjunction with data from 13,257 individuals with imputed genotype data.

## Methods

Each study had a protocol approved by its respective Institutional Review Board and participants provided informed consent. A study overview is provided in **Supplementary Figure 1**. There were two classes of data: “WGS studies” had WGS performed by the TOPMed program and, in some cases, in additional participants by the CCDG program (referred to as “WGS” studies); “Imputed studies” had array-based genotyping later imputed using the TOPMed imputation server (as described below). Some studies with WGS contributed imputed study data from additional array-based genotyped individuals.

### WGS studies

The <u>Atherosclerosis Risk in Communities Study</u> (ARIC), the <u>Cardiovascular Health Study</u> (CHS), and the <u>Framingham Heart Study Offspring Cohort</u> (FHS) included individuals who participated in the Sleep Heart Health Study (SHHS), who underwent polysomnography (PSG) between 1995 – 1998 using the Compumedics PS-2 system [^19–22^]. These samples included 1,028 EAs from ARIC; 151 AAs and 557 EAs from CHS; and 478 EAs from FHS.

The <u>Multi-Ethnic Study of Atherosclerosis</u> (MESA) is investigating the risk factors for clinical cardiovascular disease [^23^]. PSG was obtained between 2010 – 2013 using the Compumedics Somte system [^24^]. This analysis includes data from 698 EAs, 486 AAs, 456 HAs, and 229 AsAs.

The <u>Cleveland Family Study</u> (CFS) was designed to investigate the familial basis of SDB, with four visits occurring from 1990 – 2006 [^25^]. Sleep was assessed either in a clinical research center using full PSG (Compumedics E series) (visit 4); or in the latest available prior examination using an in-home sleep apnea testing device (Edentrace). Data were analyzed from 505 AAs and 485 EAs (339 AAs and 234 EAs with full PSG data).

The <u>Hispanic Community Health Study/Study of Latinos</u> (HCHS/SOL) is studying multiple health conditions in HAs [^26,27^]. Home sleep apnea testing was performed during the baseline examination (2008 – 2011) using the ARES Unicorder 5.2, a validated device including a forehead-based reflectance oximeter, a nasal pressure cannula and pressure transducer, an accelerometer, and a microphone [^28^]. 2,339 individuals provided data.

The <u>Jackson Heart Study</u> (JHS) is investigating cardiovascular disease in AAs [^29^]. An in-home sleep study was performed from 2012 – 2016 using a validated Type 3 sleep apnea testing device (Embla Embletta Gold) [^30,31^]. 575 individuals contributed data.

### Imputed genotype studies

The <u>Osteoporotic Fractures in Men Study</u> (MrOS) is a multi-center cohort study initially designed to examine the risk factors for osteoporosis, fractures, and prostate cancer in older males [^32,33^]. An ancillary study (MrOS Sleep; 2003 – 2005) focused on outcomes of sleep disturbances used PSG and nearly identical procedures as in MESA (Compumedics Safiro system) [^34^]. 2,181 EA individuals were included, with genotyping performed using the Illumina Human Omni 1 Quad v1-0 H array.

The <u>Starr County Health Studies</u> (Starr) investigates the risk factors for diabetes in Mexican-Americans [^35,36^]. An in-home sleep apnea study occurred between 2010 and 2014 using a validated instrument that records finger pulse oximetry, actigraphy, body position, and peripheral arterial tonometry (Itamar-Medical WatchPAT-200) [^37^]. 782 HA individuals were studied, using Affymetrix 6.0 genotyping data.

The <u>Western Australian Sleep Health Study</u> (WASHS) is a clinic-based study focused on the epidemiology and genetics of SDB [^38^]. PSG was obtained from 1,508 European-ancestry patients (91% referred for SDB evaluation) from 2006 – 2010 (Compumedics Series E). Genotyping was performed using the Illumina Omni 2.5 array.

Imputed genotype data were available for additional members of the TOPMed cohorts described above. Study/population combinations with fewer than 100 individuals were excluded. ARIC contributed an additional 631 EA individuals (Affymetrix 6.0; dbGaP phg000035.v1.p1). CFS contributed 225 AA and 218 EA individuals (Affymetrix 6.0; Illumina OmniExpress+Exome, Exome, and IBC). CHS contributed 365 individuals (Illumina CNV370 and IBC; phg000135.v1.p1 and phg000077.v1.p1). FHS contributed 192 EA individuals (Affymetrix 500k; phg000006.v7). HCHS/SOL contributed 7,155 HA individuals (Illumina Omni 2.5; phg000663.v1).

### Phenotype and covariate definitions

We examined several SDB measures, including specific measures of OSA: AHI (number of apneas plus hypopneas per hour of sleep, with a minimum 3% desaturation per event) and average oxyhemoglobin desaturation per apnea or hypopnea; and measures of SDB severity [^7–9^]: average and minimum SpO_2_ and the percentage of the night with SpO_2_ < 90% (Per90). Apart from WASHS, all sleep data were scored by blinded scorers at one central Sleep Reading Center with high levels of scorer reliability using well-defined procedures [^39,40^]. We adjusted for age, age^2^, sex, age × sex, body mass index (BMI), and BMI^2^ due to known age and sex effects, some of which are non-linearly associated with outcomes, and our goal of identifying obesity-independent loci. Age and BMI were obtained at the time of the sleep recording. Phenotype analyses were pooled within populations to aggregate very rare variants for testing, and therefore further adjusted for study. Cryptic relatedness and population substructure were controlled for using linear mixed models. Genomic control was applied to population-specific results (or cohort-specific imputed genotype results).

### WGS and genotyping

Sequence data were derived from the TOPMed Freeze 6a release, jointly called by the TOPMed Informatics Research Center at the University of Michigan (http://github.com/statgen/topmed_variant_calling). The methodology was described elsewhere [^17^]. In brief, WGS was performed at the Broad Institute (ARIC, FHS, MESA), Baylor College of Medicine (ARIC, CHS, HCHS/SOL), and the University of Washington (CFS, JHS). Additional ARIC and HCHS/SOL WGS funded by CCDG and performed at Baylor College of Medicine were included in the jointly-called data. TOPMed and CCDG calling pipelines have functionally equivalent outcomes despite data processing differences (as detailed in [^41^]). WGS data were merged and normalized; inferred sequence contamination was identified; and SNPs and small indels were detected (structural variants are not currently available). Lower quality variants were excluded using Mendelian consistency checks. Variants were aligned to Build 38 and annotated using snpEff 4.3t [^42^]. We excluded variants with <10× depth or >5% missingness, leaving 152.7 million polymorphic variants in 7,988 individuals with SDB phenotypes.

Genotype data were imputed using the TOPMed Imputation Server [^18^] using a Freeze 5b (Build 38) template. Forward strand checks were performed using the Strand database and the Haplotype Reference Consortium imputation preparation script (https://www.well.ox.ac.uk/~wrayner/tools/) and confirmed using Ensembl variant allele checks and internal QC performed on the server. Study-level data were imputed separately. Analyses on variants with r^2^ score > 0.5 were therefore performed separately for each study.

### Statistical analyses

Single and grouped variant analyses were performed using EMMAX and MMSKAT, both within the EPACTS suite (v3.3, https://genome.sph.umich.edu/wiki/EPACTS) [^43^]. WGS genetic relatedness matrices (GRM) were constructed using autosomal variants (MAF > 0.1%) following a comparison of EPACTS point-wise heritability estimates of the AHI using different minimal MAFs. A grid search identified optimal GRM parameters with imputed data (MAF > 0.5%, r^2^ > 0.90) using 929 ARIC individuals with imputation and WGS data. Log10 P-values using identical association test parameters had a Spearman’s ρ correlation of 0.951 between WGS and imputed data. Matrices were constructed separately for each study + population combination (due to potentially differential imputation coverage).

Gene-based group sets were constructed with a series of filters considering non-pseudogenes expressed in any GTEx v7 tissue. A variant could be assigned to one or more Ensembl genes based on SNPEff annotations [^42,44^]. We examined 5_prime_UTR_premature_start_codon_gain_variant, bidirectional_gene_fusion, conservative_inframe_deletion, conservative_inframe_insertion, disruptive_inframe_deletion, disruptive_inframe_insertion, exon_loss_variant, frameshift_variant, gene_fusion, intiator_codon_variant, missense_variant, non_canonical_start_codon, splice_acceptor_variant, splice_donor_variant, splice_region_variant, start_loss, stop_gained, stop_lost, and stop_retained_variant mutations. We also included variants located within experimentally derived promoter regions and Ensembl-derived Tarbase miRNA binding sites; and regulatory variants located within 1000 bases of a particular gene, including ChIP-seq determined transcription factor binding sites (TFBS), and Ensembl-derived CTCF, TFBS, and promoter sites [^44–46^]. Group set variants were filtered by requiring either a FATHMM-XF score > 0.5 or a CDTS < 1% constrained region score [^47,48^]. Exonic variants could alternatively have a PrimateAI score > 0.803 or a Havrilla *et al*. < 1% constrained coding region score [^49,50^].

Gene-based tests considered variants in WGS-only data (MAF < 5%). Pooled (across cohort) analyses were performed within each population in order to aggregate information on very rare variants across studies. Combined population results were obtained through meta-analysis of p-values weighted by sample size (due to potentially different MAF spectra driven by population demography). A significance level of p < 4.51 × 10^−8^ was used, reflecting a Bonferroni adjustment for all genes tested across all phenotype and population configurations.

A second set-based analysis was designed to query for TFBS annotation enrichment [^51^]. We performed 250 base-pair sliding window analyses (to improve power by aggregating additional variants beyond an approximate ChIP-seq peak width of 100 base-pairs). We filtered for variants with either a FATHMM-XF score > 0.5 or a CDTS 1% score with no MAF cut-offs and meta-analyzed MMSKAT results across the 4 populations, noting windows with p-values < 0.01. These intervals were tested for enrichment of ChIP-seq coordinates with at least 50% physical overlap for up to 437 transcription factors using ReMap 2018 v1.2 (http://tagc.univ-mrs.fr/remap/index.php?page=annotation) [^52^].

Single-variant EMMAX tests examined common variants (MAF > 0.5%). Meta-analysis across populations (and imputed genotype studies) used METAL with genomic control [^53^]. We performed bidirectional discovery and replication using the WGS and imputed samples (noting the high genomic resolution in the WGS samples and the higher sample size in the imputed data). We report results including at least 1000 individuals, discovery association p-values < 1 × 10^−5^ and replication association p-values < 0.05. Significance was defined as p < 1 × 10^−8^ in joint analyses, reflecting adjustment for five correlated phenotypes (Supplementary Table S3). We performed MetaXcan imputed GTEx gene expression analyses using joint EA results in selected tissues relevant to SDB and GIGSEA pathway analyses of MetaXcan output in whole blood (to maximize power), with empirical p-values incorporating 10,000 permutations [^54,55^].

## Results

### Study sample

A study overview is provided in **Supplementary Figure 1. Tables 1 and 2** provide a summary of the study samples and SDB traits analyzed using WGS and imputed genotypes, respectively. In total, there were 21,244 individuals (1,942 AAs; 229 AsAs; 8,341 EAs; and 10,732 HAs). Median AHI levels ranged from mildly to moderately elevated, reflecting the age range and sex distribution of each cohort. Pairwise correlations of phenotypes and covariates are provided in **Supplementary Table 3**.

**Table 1.**
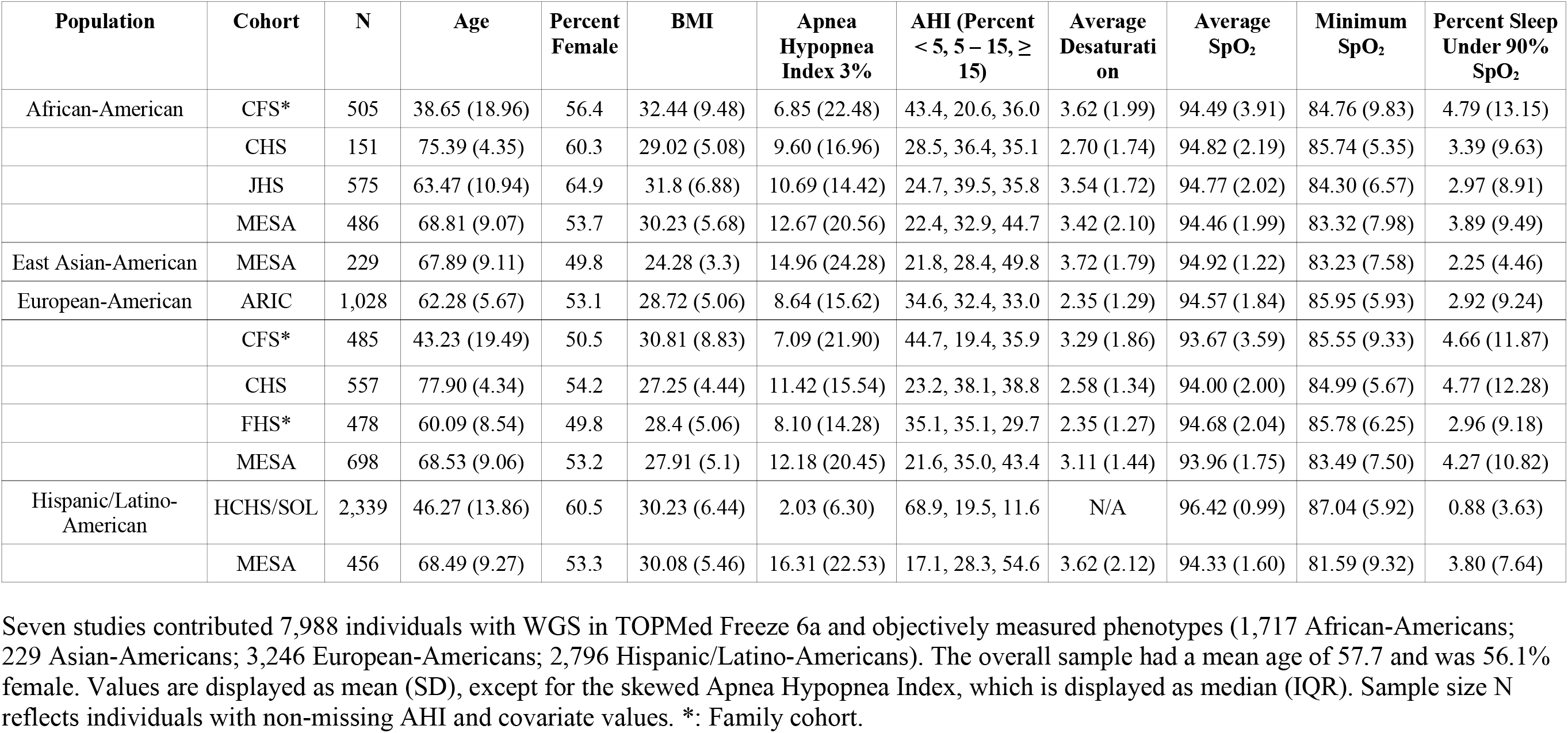
Sample description for WGS cohorts.

**Table 2.**
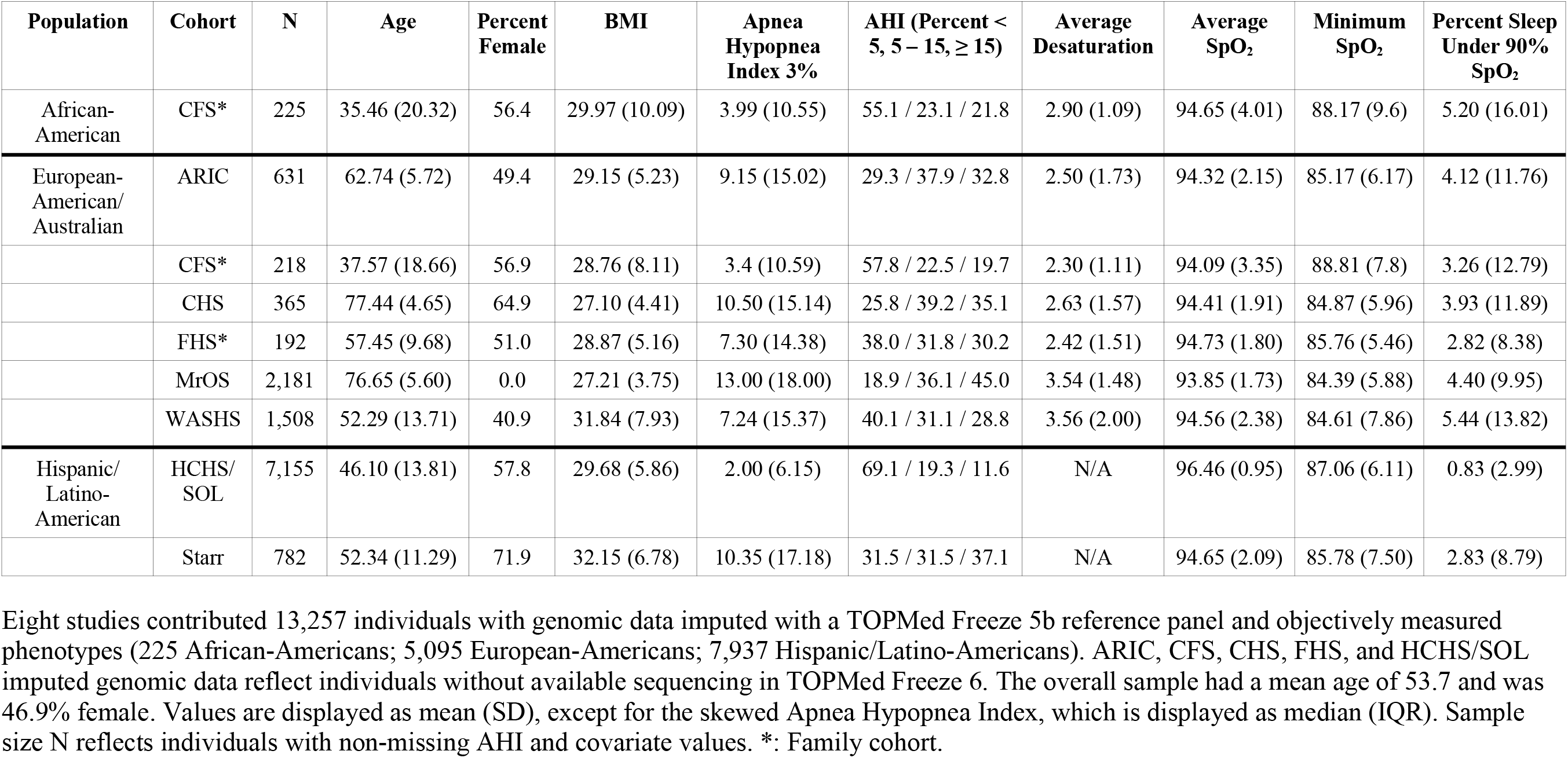
Sample description for imputed genotype cohorts.

### Gene-based results

Gene-based rare variant results are presented in **Table 3** (for meta-analyzed results across multiple populations) and in **Table 4** (for secondary population-specific results). Collectively, we identified 4 significantly associated genes (Bonferroni p < 4.51 × 10^−8^). *ARMCX3*, identified in the multiple-population analysis, is an X-linked protein-coding that was associated with average desaturation (p = 5.29 × 10^−8^). Two protein-coding genes were identified in population-specific analyses of Per90: *MRPS33* (p = 1.22 × 10^−9^) and *C16orf90* (p = 1.36 × 10^−8^). We identified 12 suggestively associated genes (p ≤ 4.22 × 10^−7^). Three genes are druggable [^56,57^]. Nominally significant results (p < 0.01) and additional details are presented in **Supplementary Tables 4 and 5**.

**Table 3.** Lead gene-based multiple population results.

| Phenotype | Sex | Gene | B38 Positions | P | Populations | N | Variants |
| --- | --- | --- | --- | --- | --- | --- | --- |
| Avg Desaturation | All | *ARMCX3* | X:101,623,082 – 101,625,765 | $3.48 \times 10^{-8}$ | AA, AsA, EA, HA | 5,222 | 8, 5, 24, 9 |
| | All | *ARMCX3-AS1* | X:101,623,082 – 101,625,153 | $3.49 \times 10^{-8}$ | AA, AsA, EA, HA | 5,222 | 7, 5, 23, 8 |
| Per90 | All | *OR5K2* | 3:98,497,633 – 98,498,634 | $2.55 \times 10^{-7}$ | AA, AsA, EA, HA | 7,986 | 4, 2, 1, 1 |
| Per90 | Females | *ZZEF1* | 17:4,004,409 – 4,144,018 | $4.22 \times 10^{-7}$ | AA, AsA, EA, HA | 4,485 | 85, 16, 87, 131 |

**Table 4.**
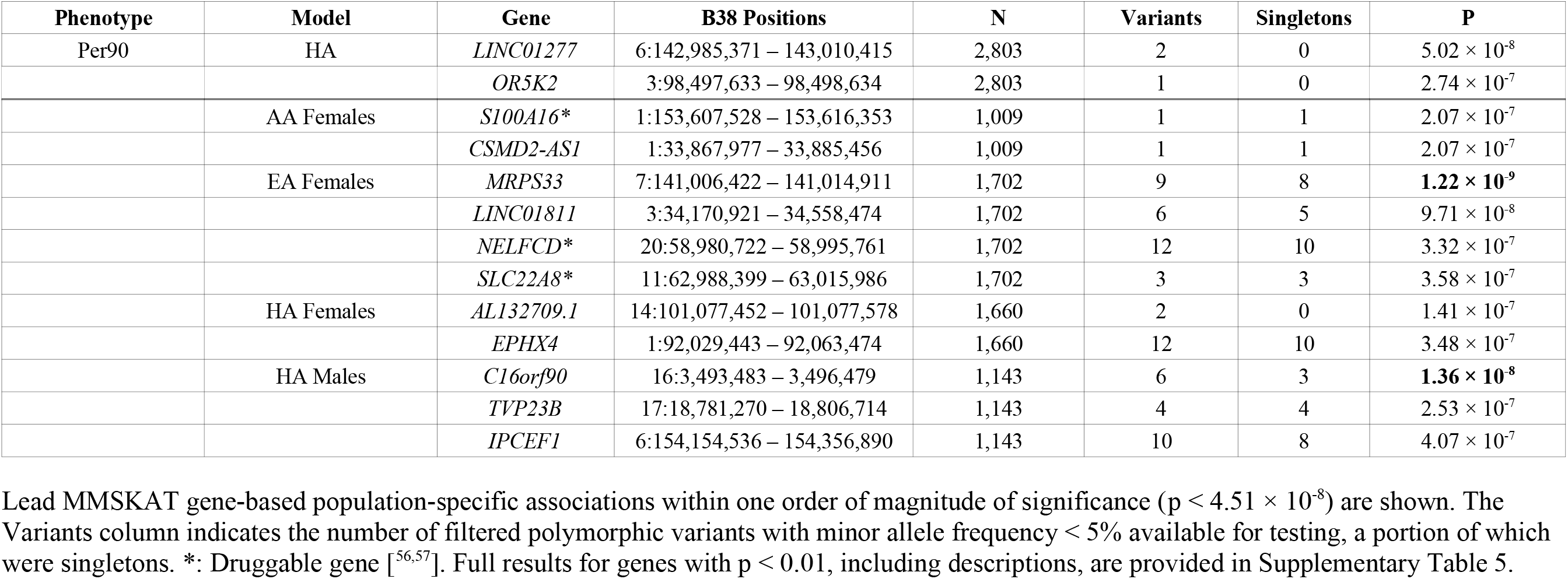
Lead gene-based population-specific results.

### Single-variant results

We identified four genome-level significant loci in single-variant analyses (MAF > 0.5%; p < 1.0 × 10^−8^; **Table 5**). In multiple-population analyses, the 2q12 locus (rs77375846; *IL18RAP*) was associated with average event desaturation in a multiple-population analysis (combined p = 1.57 × 10^−9^) and minimum SpO_2_ (consistent with a previous report [^14^]). Two novel population-specific loci were identified. The 8p12 locus (rs35447033, *NRG1*) was associated with AHI in EAs (combined p = 3.02 × 10^−9^, **Figure 1**). The 5p13 locus (rs28777; *SLC45A2*) was associated with average SpO_2_ in EAs (combined p = 8.08 × 10^−10^, **Figure 2**). In HAs, the 1q32 locus (rs116133558; *ATP2B4*) was associated with Per90 (combined p = 3.51 × 10^−10^) and with average SpO_2_ (as previously identified [^11^]). Twelve additional regions were suggestively associated (p < 1.0 × 10^−7^). **Supplementary Table 6** provides additional context for all variants in these loci (p < 1.0 × 10^−7^), including imputation quality, significant eQTLs, and overlap with epigenetic regions [^58–61^]. Manhattan and QQ plots corresponding to the significant associations are provided in **Supplementary Figures 2 – 5**.

**Table 5.** Lead single-variant analysis results.

| Region | Phenotype | Model | SNP | WGS /<br>Imputed N | CAF | WGS Beta<br>(SE) | WGS P | Imputed Beta<br>(SE) | Imputed P | Combined<br>Beta (SE) | Combined<br>P |
| --- | --- | --- | --- | --- | --- | --- | --- | --- | --- | --- | --- |
| *2q12.1: IL18RAP* | Avg<br>desaturation | All | rs77375846 C | 4995 / 4838 | 0.028 – 0.129 | -0.152 (0.049) | $1.87 \times 10^{-3}$ | -0.264 (0.049) | $5.97 \times 10^{-8}$ | -0.208 (0.035) | **$1.57 \times 10^{-9}$** |
| *2q33.3: PPIAP68* | Avg<br>desaturation | All | rs60132122 T | 5222 / 4838 | 0.308 – 0.637 | 0.062 (0.031) | 0.043 | 0.195 (0.034) | $6.26 \times 10^{-9}$ | 0.122 (0.023) | $6.49 \times 10^{-8}$ |
| *11q12.2: MS4A15* | Avg SpO<sub>2</sub> | All | rs4939452 C | 7929 / 13197 | 0.347 – 0.524 | 0.066 (0.023) | $4.34 \times 10^{-3}$ | 0.063 (0.014) | $3.29 \times 10^{-6}$ | 0.064 (0.012) | $4.87 \times 10^{-8}$ |
| *18q12.3:*<br>*LINC00907* | Avg SpO<sub>2</sub> | All | rs187860354 G | 4500 / 7391 | 0.006 – 0.022 | 0.442 (0.146) | $2.36 \times 10^{-3}$ | 0.432 (0.097) | $8.53 \times 10^{-6}$ | 0.436 (0.081) | $7.04 \times 10^{-8}$ |
| *2q12.1: IL18RAP* | Min SpO<sub>2</sub> | All | rs138895820 G | 7705 / 13194 | 0.025 – 0.131 | 0.510 (0.184) | $5.58 \times 10^{-3}$ | 0.654 (0.128) | $3.36 \times 10^{-7}$ | 0.607 (0.105) | **$7.93 \times 10^{-9}$** |
| *10p12.31: NEBL* | Min SpO<sub>2</sub> | Females | rs11453507 CA | 4450 / 6202 | 0.138 – 0.514 | 0.651 (0.140) | $3.34 \times 10^{-6}$ | 0.338 (0.102) | $8.63 \times 10^{-4}$ | 0.446 (0.082) | $5.73 \times 10^{-8}$ |
| *12q21.2:*<br>*LINC024064* | Min SpO<sub>2</sub> | Females | rs2176909 T | 4450 / 6202 | 0.724 – 0.930 | 0.828 (0.157) | $1.38 \times 10^{-7}$ | 0.319 (0.116) | $5.77 \times 10^{-3}$ | 0.498 (0.093) | $9.06 \times 10^{-8}$ |
| *5p13.3: C5orf22* | AHI | Males | rs10940956 A | 3502 / 7043 | 0.470 – 0.759 | 0.930 (0.422) | $2.74 \times 10^{-2}$ | 1.430 (0.269) | $1.09 \times 10^{-7}$ | 1.285 (0.227) | $1.48 \times 10^{-8}$ |
| *9p22.1:*<br>*DENND4C* | AHI | AA | rs111654000 A | 1717 / 225 | 0.016 – 0.018 | -11.240<br>(2.268) | $7.18 \times 10^{-7}$ | -18.110 (6.724) | $7.07 \times 10^{-3}$ | -11.942<br>(2.149) | $2.74 \times 10^{-8}$ |
| *1q31.2:*<br>*AL954650.1* | AHI | AA | chr1:191965014_G/<br>A A | 1717 / 225 | 0.286 – 0.301 | 3.078 (0.641) | $1.56 \times 10^{-6}$ | 5.080 (1.759) | $3.88 \times 10^{-3}$ | 3.313 (0.602) | $3.75 \times 10^{-8}$ |
| *8p12:*<br>*AC068672.1,*<br>*NRG1* | AHI | EA | rs35447033 T | 3246 / 5095 | 0.060 – 0.094 | 2.247 (0.621) | $2.95 \times 10^{-4}$ | 2.453 (0.521) | $2.54 \times 10^{-6}$ | 2.368 (0.399) | **$3.02 \times 10^{-9}$** |
| *5p13.2: SLC45A2* | Avg SpO<sub>2</sub> | EA | rs28777 A | 3201 / 5024 | 0.885 – 0.969 | -0.526 (0.133) | $8.00 \times 10^{-5}$ | -0.454 (0.096) | $2.23 \times 10^{-6}$ | -0.478 (0.078) | **$8.08 \times 10^{-10}$** |
| *1q32.1: ATP2B4* | Avg SpO<sub>2</sub> | HA | rs116133558 T | 2803 / 7956 | 0.006 – 0.014 | 0.371 (0.120) | $2.08 \times 10^{-3}$ | 0.294 (0.062) | $2.15 \times 10^{-6}$ | 0.310 (0.055) | $1.88 \times 10^{-8}$ |
| *1q23.3: intergenic*<br>*(RNU6-755P)* | Min SpO<sub>2</sub> | HA | rs140743827 A | 2803, 7174 | 0.017 – 0.020 | -1.502 (0.593) | $1.13 \times 10^{-2}$ | -1.770 (0.367) | $1.42 \times 10^{-6}$ | -1.696 (0.312) | $5.51 \times 10^{-8}$ |
| *1q32.1: ATP2B4* | Per90 | HA | rs116133558 T | 2803, 7956 | 0.006 – 0.014 | -1.005 (0.450) | $2.54 \times 10^{-2}$ | -1.218 (0.207) | $4.15 \times 10^{-9}$ | -1.181 (0.188) | **$3.51 \times 10^{-10}$** |
| *11p11.2:*<br>*intergenic*<br>*(AC104010.1)* | Avg SpO<sub>2</sub> | HA males | chr11:44652095_T<br>C/T T | 1143 / 3024 | 0.007 – 0.008 | 0.686 (0.248) | $5.65 \times 10^{-3}$ | 0.710 (0.154) | $3.83 \times 10^{-6}$ | 0.703 (0.131) | $7.25 \times 10^{-8}$ |
| *10q22.1: HK1* | Min SpO<sub>2</sub> | EA males | rs17476364 C | 1523 / 3650 | 0.072 – 0.115 | 1.215 (0.392) | $1.94 \times 10^{-3}$ | 1.099 (0.235) | $2.81 \times 10^{-6}$ | 1.129 (0.201) | $2.01 \times 10^{-8}$ |
| *8q23.2: KCNV1* | Min SpO<sub>2</sub> | EA males | rs58365105 A | 1523, 3650 | 0.007 – 0.026 | -2.878 (0.864) | $8.65 \times 10^{-4}$ | -2.406 (0.540) | $8.36 \times 10^{-6}$ | -2.539 (0.458) | $2.96 \times 10^{-8}$ |
| *2q35:*<br>*AC019211.1* | Per90 | EA males | chr2:220369683_G/<br>A A | 1540, 187 | 0.005 – 0.006 | 12.280 (2.431) | $4.38 \times 10^{-7}$ | 17.505 (7.989) | $2.85 \times 10^{-2}$ | 12.723 (2.326) | $4.48 \times 10^{-8}$ |

**Figure 1.**
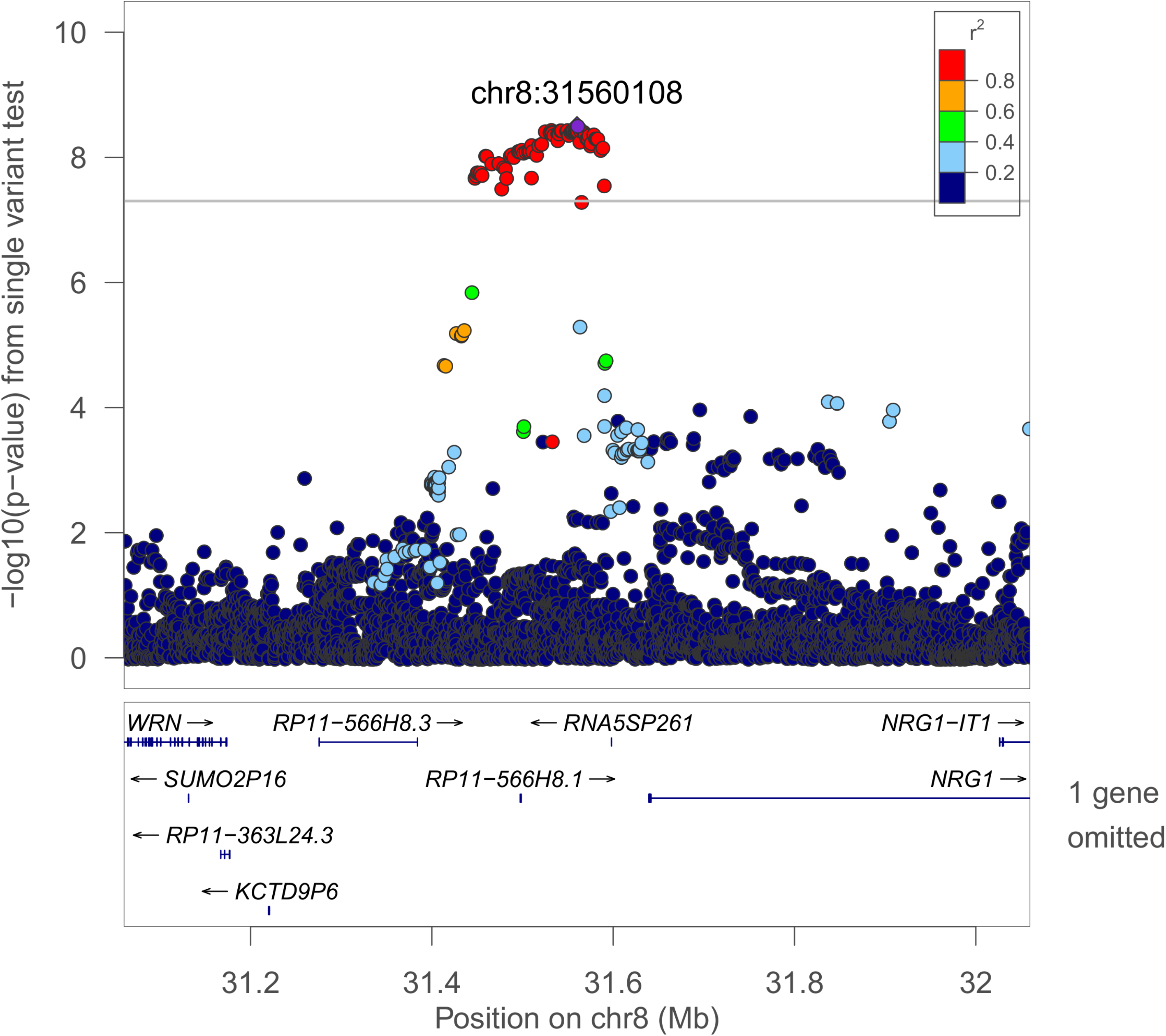
Regional plot of the rs35447033 association with AHI in European-ancestry individuals. Joint WGS and imputed results are shown, using Build 38 coordinates.

**Figure 2.**
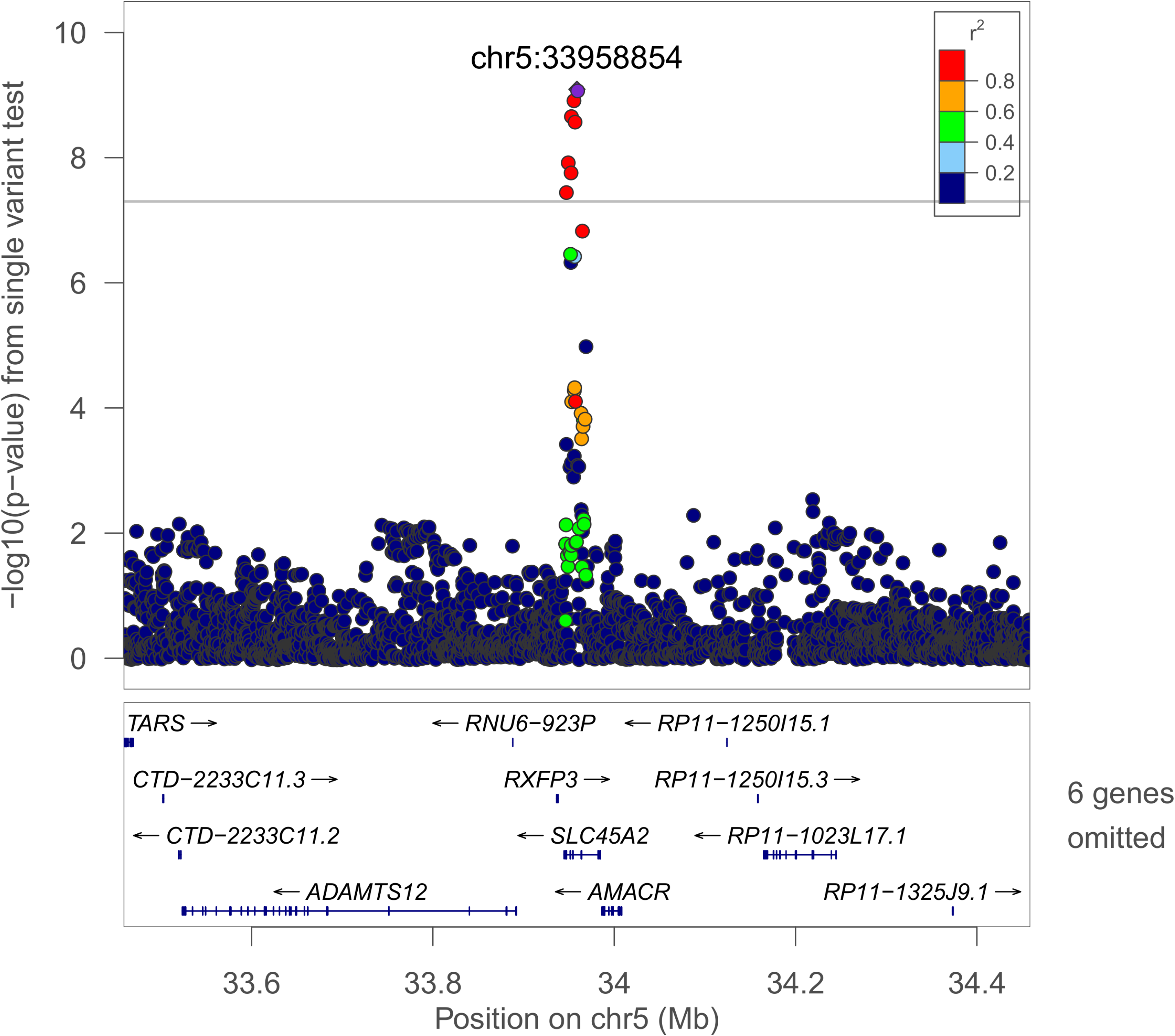
Regional plot of the rs28777 association with Average SpO_2_ in European-ancestry individuals. Joint WGS and imputed results are shown, using Build 38 coordinates.

### MetaXcan imputed gene expression and GIGSEA pathway analyses

We used joint WGS and imputed EA results to impute associations with gene expression levels using a MetaXcan framework for 6 tissues (subcutaneous and visceral omentum adipose, lung, monocytes, skeletal muscle, and whole blood). No individual tests reached Bonferroni significance (p < 2.60 × 10^−7^; **Supplementary Table 7**). Genes that were observed in the top 10 results across the varied analyses (**Supplementary Table 8**) included *ZNF83* (15 instances) and *CHRNE* (13 instances).

Whole blood MetaXcan results (with the largest sample size) were further evaluated in GIGSEA-based pathway analyses. KEGG pathway results are shown in **Supplementary Table 9**. The most significantly associated pathway was KEGG_STEROID_HORMONE_BIOSYNTHESIS (average SpO_2_ empirical p-value = 7.00 × 10^−4^). KEGG_RIG_I_LIKE_RECEPTOR_SIGNALING_PATHWAY was observed in the top 10 results for 4 of the 5 phenotypes. Gene-centric transcription factor binding site (TFBS) enrichment analysis results are presented in **Supplementary Table 10**. V$PEA3_Q6 (*ETV4*) was the most significantly associated TFBS (average desaturation empirical p-value = 3.00 × 10^−4^) and was the strongest association for AHI and minimum SpO_2_ (empirical p-values 0.002 and 0.001, respectively). The most significant miRNA binding site enrichment analysis association was GCATTTG,MIR-105 (average SpO_2_ p = 0.002; **Supplementary Table 11**). AGGCACT,MIR-515-3P (the strongest AHI association, p = 0.009) was observed in the top ten results for four phenotypes.

### ChIP-seq transcription factor binding site interval enrichment

We performed a sliding window analysis to examine enriched intervals containing ChIP-seq derived coordinates for up to 437 transcription factors (**Table 6, Supplementary Table 12**). *FOXP2* TFBS were consistently the most enriched for all phenotypes. Other notable transcription factors in the top 5 included *EGR1, KDM4B, KDM6B*, and *TP63. KDM4B* and *KDM6B* are druggable [^56,57^]. Leading sliding window results are provided in **Supplementary Table 13**.

**Table 6.** Transcription factor binding site interval enrichment results.

| Phenotype | Transcription Factor | # Observed Overlap | # Expected Overlap | -log<sub>10</sub> (E-value) |
| --- | --- | --- | --- | --- |
| AHI | *FOXP2* | 588 | 36.20 | 473.99 |
|  | *KDM6B* | 630 | 51.58 | 435.29 |
|  | *THAP1* | 505 | 31.89 | 402.07 |
|  | *KLF9* | 745 | 91.81 | 395.52 |
|  | *TP63* | 997 | 182.22 | 383.85 |
| Average Desaturation | *FOXP2* | 493 | 22.32 | 460.00 |
|  | *THAP1* | 439 | 19.55 | 412.76 |
|  | *UBTF* | 489 | 28.20 | 407.50 |
|  | *TP63* | 788 | 109.36 | 382.89 |
|  | *KDM6B* | 482 | 30.98 | 380.39 |
| Average SpO<sub>2</sub> | *FOXP2* | 582 | 35.87 | 468.89 |
|  | *KDM6B* | 613 | 51.21 | 418.65 |
|  | *EGR1* | 664 | 66.76 | 404.83 |
|  | *UBTF* | 574 | 46.35 | 399.91 |
|  | *KDM4B* | 489 | 29.56 | 398.10 |
| Min SpO<sub>2</sub> | *FOXP2* | 561 | 35.57 | 445.57 |
|  | *THAP1* | 515 | 31.32 | 417.89 |
|  | *KDM6B* | 569 | 50.87 | 373.41 |
|  | *UBTF* | 536 | 45.99 | 360.56 |
|  | *EGR1* | 602 | 66.25 | 346.03 |
| Per90 | *FOXP2* | 689 | 39.05 | 578.42 |
|  | *KDM6B* | 739 | 54.79 | 539.69 |
|  | *TP63* | 1199 | 193.28 | 515.44 |
|  | *THAP1* | 607 | 34.47 | 509.33 |
|  | *EGR1* | 786 | 72.09 | 507.27 |

## Discussion

Sleep-disordered breathing (SDB) is associated with increased risk of a wide range of disorders, including atrial fibrillation, cancer, cognitive impairment, diabetes, liver, and interstitial lung diseases, as well as premature mortality [^3,4,62–67^]. Treatment options, however, are limited by a lack of knowledge of molecular pathways, including those that may be “druggable”. Recent analyses of SDB traits have focused on common variants and identified several preliminary genome-level significant associations using GWAS, admixture mapping, and linkage approaches [^11–15^], but did not address gene-based or rare variant effects. Ten studies and over 21,000 individuals of multiple ancestries with WGS data at unprecedented resolution from the NHLBI TOPMed program combined with densely imputed data from other sources contributed to these results. We identified several variant, gene-based, and pathway-level associations. Analyses adjusted for obesity, a major SDB risk factor, identified loci and genes implicated in pulmonary, inflammatory, and craniofacial pathways. Some associations were population-specific, while others were sex-specific, consistent with population differences and strong sex differences for SDB [^24,68–70^]. Notably, across multiple ancestral groups, we identified a set-based rare-variant association (p = 3.48 × 10^−8^) on chromosome X with *ARMCX3*.

### Gene-based results

Across multiple populations, *ARMCX3 (ALEX3*) and the RNA anti-sense gene *ARMCX3-AS1* were associated with apnea-hypopnea triggered intermittent hypoxia. *ARMCX3* regulates mitochondrial aggregation and trafficking in multiple tissues and facilitates neuronal survival and axon regeneration [^71–73^]. Wnt signaling regulates reactive oxygen species (ROS) generation and *ARMCX3*-associated mitochondrial aggregation [^72,74^]. Potential mechanisms for further study include sensitized carotid body chemoreflexes, interaction with inflammatory mechanisms, and neuronal dysfunction within respiratory centers. Sleep apnea and reduced ventilatory drive are enriched in individuals with a primary mitochondrial disorder [^75^]. Mitochondria are an important source of ROS, which modulate the acute hypoxic ventilatory response. Mitochondria impact *HIF1A* signaling and may contribute to oxygen sensing [^76–79^]. ROS are required for intermittent hypoxia-induced respiratory long-term facilitation [^80,81^]. These effects may mitigate the level of hypoxia resulting from recurrent apneas, or conversely, lead to ventilatory instability, promoting apnea occurrence. Mitochondrial ROS also activate the NLRP3 inflammasome in multiple pulmonary diseases, consistent with an inflammation model that includes our IL18-pathway and *HK1* results, ROS-related proinflammatory responses to lung capillary pressure, and evidence of alveolar epithelial injury/SDB interactions [^14,82–87^]. Our findings suggest value in investigating the mechanisms by which *ARMCX3* predisposes to SDB, and whether these associations are mediated by neuronal dysfunction and/or ROS and carotid body sensitization, and interact with the inflammasome.

Additional genes were significantly associated in population-specific analyses, including the mitochondrial ribosomal gene *MRPS33*. Mitoribosomes are responsible for expression of the 13 essential components of the oxidative phosphorylation system, and a majority of the small subunit proteins have been implicated in disease [^88^]. The expression of several small and large subunit proteins are altered in a hypoxic environment [^89^]. *MRPS33* expression varies with oxygen treatment in COPD [^90^].

### Single-variant results

We identified four common frequency associated loci, including multiple-population associations with the *IL18RAP* region. The *IL18RAP* region has been associated with minimum SpO_2_ [^14^], and here we further identify an association with average event desaturation, highlighting a role in an OSA-specific trait. Multiple variants in this region are also GTEx eQTL variants for both interleukin-18 receptor subunits *IL18RAP* and *IL18R1* (Supplementary Table 6) and experimental studies support a role for *IL18* signaling in mediating this association, possibly through effects of pulmonary inflammation on gas exchange (reviewed in [^14^]).

We identified three population-specific loci, including two novel associations in individuals of European ancestry (Figures 1 and 2). 65 variants in the *NRG1* region were associated with the AHI (p < 1.0 × 10^−8^, Supplementary Table 6). This region was suggestively associated with sleep apnea in a Korean population [^91^], however the lead signals appear to be independent (rs10097555 Korean p = 2.6 × 10^−6^, EA p = 0.91). *NRG1* is associated with lung development and acute lung injury, and mediates inflammasome-induced alveolar cell permeability [^87,92–95^]. NRG1 promotes accumulation of HIF1A and has increased expression in vascular smooth muscle cells following exposure to intermittent hypoxia [^96,97^]. The lead *SLC45A2* region variant rs28777 (average SpO_2_ p = 8.08 × 10^−10^) has been associated with multiple traits and is in a splicing regulatory element with extreme population differentiation [^98^]. An association in the *ATP2B4* region with average SpO_2_ in HAs [^11^] has been extended to a second hypoxemia trait at the same variant (Per90 p = 3.31 × 10^−10^). This gene is the main cellular membrane calcium pump in erythrocytes and also regulates vascular tone [^99,100^].

### Pathway analyses

Several gene pathways were identified in EA individuals using imputed gene expression in whole blood (Supplementary Table 9). KEGG_RIG_I_LIKE_RECEPTOR_SIGNALING_PATHWAY (retinoic acid-inducible gene I-like) was the most commonly observed, occurring in the top 10 results for 4 of the 5 phenotypes. This pathway initiates the immune response to RNA virus infection [^101^], consistent with a role for inflammation at the *NRG1* and *IL18RAP* loci. Steroid hormone biosynthesis (the most significantly associated pathway), PPAR signaling, and metabolism (via ‘starch and sucrose metabolism’) suggest the importance of biological pathways modulating energy homeostasis and balance and metabolic function [^102^]. In the gene-centric GIGSEA TFBS analysis, V$PEA3_Q6 (*ETV4*) was the lead association for three phenotypes. *ETV4* influences branching in the developing lung and regulates hypoxia-inducible factor signaling [^103,104^], a major mechanism influencing ventilatory control.

### Transcription factor binding site enrichment

Several transcription factors were identified through interval enrichment of observed TFBS across the genome (Table 6). *FOXP2* was consistently the most enriched transcription factor and is known to regulate gene expression in epithelial lung tissue and response to lung injury through an inflammatory mechanism [^105,106^]. *FOXP2* is also expressed in brainstem respiratory areas including the pre-Bötzinger complex (which is essential for respiratory rhythmogenesis) and impacts airway morphology [^107,108^]. Two lysine demethylases (*KDM4B* and *KDM6B*) were also identified. *KDM6B (JMJD3*) is required for a functional pre-Bötzinger complex [^109,110^] and reduced KDM6B protein expression was reported in hypoxic OSA patients [^111^]. *Kdm6b* also plays roles in immune function and lung development [^112–114^]. Drosophila *Kdm4b* knock-outs have increased sleep [^115^]. *KDM4B (JMJD2B*) and *KDM6B* are both members of the JmjC protein domain family and are regulated by *HIF1A*, require oxygen as a cofactor and act as oxygen sensors for chromatin in hypoxia [^116,117^]. *EGR1* mediates hypoxia-induced pulmonary fibrosis [^118^]. *TP63* is associated with cleft palate in *Tp63* deficient mice, which is associated with an increased prevalence of OSA [^119,120^], suggesting that its relationship to OSA may be through pathways influencing craniofacial development. Among the leading 250-base pair sliding window results (Supplementary Table 13), 4:105708751-105709001 (Per90 HA p = 2.72 × 10^−9^) is of note due to regional associations with lung function and expression in human lung [^121^].

### Strengths and weaknesses

This study is the first genome-wide analysis of objectively measured SDB traits using deep sequencing. Together with improved imputation quality, the TOPMed resource has enabled unprecedented genetic resolution. We examined clinically relevant phenotypes measured using rigorous methodology [^5,7–10^]. We analyzed data from 10 studies of individuals from four population groups that used different ascertainment strategies, which may potentially improve the generalization of our results. While this analysis is among the largest performed for SDB traits to date, our moderate sample size has lower power to detect weaker associations, and data were not available to replicate these first rare variant associations. While there are multiple lines of evidence in the literature to support our findings, additional experimental followup analyses are required.

### Conclusion

We have identified the first rare-variant and additional common-variant associations at genome-level significance with objectively measured SDB traits in humans. The results point to biologically relevant pathways for further study, including a novel X-linked association (*ARCMX3*), and a number of associations in genes that modulate lung development, inflammation, respiratory rhythmogenesis and *HIF1A*-mediated hypoxic-response pathways. These associations will motivate future sample collection and follow-up in cell-line and animal validation studies, with potential therapeutic benefit for sleep-disordered breathing and related comorbidities.

## Supplementary Data

Supplementary data include 5 figures and 13 tables.

## Acknowledgements

The authors wish to thank the participants and study staff of all of our cohorts for their important contributions.

Whole genome sequencing (WGS) for the Trans-Omics in Precision Medicine (TOPMed) program was supported by the National Heart, Lung and Blood Institute (NHLBI). WGS for “NHLBI TOPMed: Trans-Omics for Precision Medicine Whole Genome Sequencing Project: ARIC (phs001211.v1.p1) was performed at Baylor College of Medicine Human Genome Sequencing Center (HHSN268201500015C and 3U54HG003273-12S2) and the Broad Institute of MIT and Harvard (3R01HL092577-06S1). WGS for “NHLBI TOPMed: The Cleveland Family Study (WGS) (phs000954.v2.p1) was performed at University of Washington Northwest Genomics Center (3R01HL098433-05S1). WGS for “NHLBI TOPMed: Cardiovascular Health Study (phs001368.v1.p1) was performed at Baylor College of Medicine Human Genome Sequencing Center (HHSN268201500015C). WGS for “NHLBI TOPMed: Whole Genome Sequencing and Related Phenotypes in the Framingham Heart Study (phs000974.v3.p2) was performed at the Broad Institute of MIT and Harvard (3R01HL092577-06S1). WGS for “NHLBI TOPMed: Hispanic Community Health Study/Study of Latinos (HCHS/SOL) (phs001395) was performed at the Baylor College of Medicine Human Genome Sequencing Center (HHSN268201500015C and 3U54HG003273-12S2). WGS for “NHLBI TOPMed: The Jackson Heart Study (phs000964.v3.p1) was performed at University of Washington Northwest Genomics Center (HHSN268201100037C). WGS for “NHLBI TOPMed: NHLBI TOPMed: MESA (phs001416.v1.p1) was performed at the Broad Institute of MIT and Harvard (3U54HG003067-13S1). Centralized read mapping and genotype calling, along with variant quality metrics and filtering were provided by the TOPMed Informatics Research Center (3R01HL-117626-02S1). Phenotype harmonization, data management, sample-identity QC, and general study coordination, were provided by the TOPMed Data Coordinating Center (3R01HL-120393-02S1). We gratefully acknowledge the studies and participants who provided biological samples and data for TOPMed.

The Genome Sequencing Program (GSP) was funded by the National Human Genome Research Institute (NHGRI), the National Heart, Lung, and Blood Institute (NHLBI), and the National Eye Institute (NEI). The GSP Coordinating Center (U24 HG008956) contributed to cross-program scientific initiatives and provided logistical and general study coordination. The Centers for Common Disease Genomics (CCDG) program was supported by NHGRI and NHLBI, and CCDG-funded whole genome sequencing of the ARIC and HCHS/SOL studies was performed at the Baylor College of Medicine Human Genome Sequencing Center (UM1 HG008898 and R01HL059367).

Brian Cade is supported by grants from the National Institutes of Health [K01-HL135405-01, R01-HL113338-04, R35-HL135818-01] and the American Thoracic Society Foundation (http://foundation.thoracic.org). Susan Redline is partially supported by grants from the National Institutes of Health [R35-HL135818-01, R01-HL113338-04]. Sanjay Patel has had grant support through his institution from the ResMed Foundation, the American Sleep Medicine Foundation, Bayer Pharmaceuticals, and Philips Respironics. James Wilson is supported by U54GM115428 from the National Institute of General Medical Sciences.

The Atherosclerosis Risk in Communities (ARIC) study has been funded in whole or in part with Federal funds from the National Heart, Lung, and Blood Institute, National Institutes of Health, Department of Health and Human Services (contract numbers HHSN268201700001I, HHSN268201700002I, HHSN268201700003I, HHSN268201700004I and HHSN268201700005I), R01HL087641, R01HL059367 and R01HL086694; National Human Genome Research Institute contract U01HG004402; and National Institutes of Health contract HHSN268200625226C. The authors thank the staff and participants of the ARIC study for their important contributions. Infrastructure was partly supported by Grant Number UL1RR025005, a component of the National Institutes of Health and NIH Roadmap for Medical Research.

This Cardiovascular Health Study (CHS) research was supported by NHLBI contracts HHSN268201200036C, HHSN268200800007C, HHSN268200960009C, HHSN268201800001C N01HC55222, N01HC85079, N01HC85080, N01HC85081, N01HC85082, N01HC85083, N01HC85086; and NHLBI grants U01HL080295, U01HL130114, R01HL087652, R01HL105756, R01HL103612, R01HL085251, and R01HL120393 with additional contribution from the National Institute of Neurological Disorders and Stroke (NINDS). Additional support was provided through R01AG023629 from the National Institute on Aging (NIA). A full list of principal CHS investigators and institutions can be found at CHS-NHLBI.org.

The Cleveland Family Study has been supported by National Institutes of Health grants [R01-HL046380, KL2-RR024990, R35-HL135818, and R01-HL113338].

The Framingham Heart Study (FHS) has been supported by contracts N01-HC-25195 and HHSN268201500001I and grant R01 HL092577. The Framingham Heart Study thanks the study participants and the multitude of investigators who over its 70 year history continue to contribute so much to further our knowledge of heart, lung, blood and sleep disorders and associated traits.

The Hispanic Community Health Study/Study of Latinos was carried out as a collaborative study supported by contracts from the NHLBI to the University of North Carolina (HHSN268201300001I / N01-HC65233), University of Miami (HHSN268201300004I / N01-HC65234), Albert Einstein College of Medicine (HHSN268201300002I / N01-HC65235), University of Illinois at Chicago (HHSN268201300003I), Northwestern University (N01-HC65236), and San Diego State University (HHSN268201300005I / N01-HC65237). The following Institutes/Centers/Offices contribute to the HCHS/SOL through a transfer of funds to the NHLBI: National Institute on Minority Health and Health Disparities, National Institute on Deafness and Other Communication Disorders, National Institute of Dental and Craniofacial Research, National Institute of Diabetes and Digestive and Kidney Diseases, National Institute of Neurological Disorders and Stroke, NIH Institution-Office of Dietary Supplements. The Genetic Analysis Center at Washington University was supported by NHLBI and NIDCR contracts (HHSN268201300005C AM03 and MOD03). The views expressed in this manuscript are those of the authors and do not necessarily represent the views of the National Heart, Lung, and Blood Institute; the National Institutes of Health; or the U.S. Department of Health and Human Services. This manuscript was not approved by the HCHS/SOL publications committee. The authors thank the staff and participants of HCHS/SOL for their important contributions. Investigator’s website - http://www.cscc.unc.edu/hchs/

The Jackson Heart Study (JHS) is supported and conducted in collaboration with Jackson State University (HHSN268201800013I), Tougaloo College (HHSN268201800014I), the Mississippi State Department of Health (HHSN268201800015I/HHSN26800001) and the University of Mississippi Medical Center (HHSN268201800010I, HHSN268201800011I and HHSN268201800012I) contracts from the National Heart, Lung, and Blood Institute (NHLBI) and the National Institute for Minority Health and Health Disparities (NIMHD). The authors also wish to thank the staff and participants of the JHS.

MESA and the MESA SHARe project are conducted and supported by the National Heart, Lung, and Blood Institute (NHLBI) in collaboration with MESA investigators. Support for MESA is provided by contracts HHSN268201500003I, N01-HC-95159, N01-HC-95160, N01-HC-95161, N01-HC-95162, N01-HC-95163, N01-HC-95164, N01-HC-95165, N01-HC-95166, N01-HC-95167, N01-HC-95168, N01-HC-95169, UL1-TR-000040, UL1-TR-001079, UL1-TR-001420. MESA Family is conducted and supported by the National Heart, Lung, and Blood Institute (NHLBI) in collaboration with MESA investigators. Support is provided by grants and contracts R01HL071051, R01HL071205, R01HL071250, R01HL071251, R01HL071258, R01HL071259, and by the National Center for Research Resources, Grant UL1RR033176. The provision of genotyping data was supported in part by the National Center for Advancing Translational Sciences, CTSI grant UL1TR001881, and the National Institute of Diabetes and Digestive and Kidney Disease Diabetes Research Center (DRC) grant DK063491 to the Southern California Diabetes Endocrinology Research Center.

The Osteoporotic Fractures in Men (MrOS) Study is supported by NIH funding. The following institutes provide support: the National Institute on Aging (NIA), the National Institute of Arthritis and Musculoskeletal and Skin Diseases (NIAMS), NCATS, and NIH Roadmap for Medical Research under the following grant numbers: U01 AG027810, U01 AG042124, U01 AG042139, U01 AG042140, U01 AG042143, U01 AG042145, U01 AG042168, U01 AR066160, and UL1 TR000128. The NHLBI provides funding for the MrOS Sleep ancillary study “Outcomes of Sleep Disorders in Older Men” under the following grant numbers: R01 HL071194, R01 HL070848, R01 HL070847, R01 HL070842, R01 HL070841, R01 HL070837, R01 HL070838, and R01 HL070839. The NIAMS provides funding for the MrOS ancillary study ‘Replication of candidate gene associations and bone strength phenotype in MrOS’ under the grant number R01 AR051124. The NIAMS provides funding for the MrOS ancillary study ‘GWAS in MrOS and SOF’ under the grant number RC2 AR058973.

The Starr County Health Studies is supported in part by grants R01 DK073541, U01 DK085501, R01 AI085014, and R01 HL102830 from the National Institutes of Health, and funds from the University of Texas Health Science Center at Houston. We thank the field staff in Starr County for their careful collection of these data and are especially grateful to the participants who so graciously cooperated and gave of their time.

Funding for the Western Australian Sleep Health Study was obtained from the Sir Charles Gairdner and Hollywood Private Hospital Research Foundations, the Western Australian Sleep Disorders Research Institute, and the Centre for Genetic Epidemiology and Biostatistics at the University of Western Australia. Funding for the GWAS genotyping obtained from the Ontario Institute for Cancer Research and a McLaughlin Centre Accelerator Grant from the University of Toronto.

## Author Contributions

Conception and design: B.E.C., G.R.A., E.A.B., A.C., L.A.C., R.C.K., D.A.N., K.E.N., B.M.P., J.I.R., S.S.R., R.P.T., R.S.V., J.G.W., S.R.

Data acquisition: B.E.C., J.L., T.S., M.Z., H.C., S.A.G., D.J.G., J.M.L., J.L., X.L., H.M., S.R.P., S.M.P., R.S., N.A.S., H.W., X.Z., D.S.E., C.L.H., D.R.H., S.M., L.J.P., K.L.S., G.J.T., G.R.A., D.A.N., S.R.

Analysis: B.E.C., J.L.

Interpretation, draft and review, and final approval: all authors.

B.E.C. and S.R. had full access to the study data and take responsibility for the integrity of the data and accuracy of analyses.

## Competing Interests

The authors disclose the following industry funding, which they believe is unrelated to this study. SRP has had grant support through his institution from the ResMed Foundation, Bayer Pharmaceuticals, and Philips Respironics. BMP serves on the Steering Committee of the Yale Open Data Access Project, funded by Johnson & Johnson. KLS has grant funding from Merck.

## References

1. Marin, J. M., Carrizo, S. J., Vicente, E. & Agusti, A. G. N. Long-term cardiovascular outcomes in men with obstructive sleep apnoea-hypopnoea with or without treatment with continuous positive airway pressure: an observational study. Lancet 365, 1046–53 (2005).

2. Strausz, S. et al. Obstructive sleep apnoea and the risk for coronary heart disease and type 2 diabetes: a longitudinal population-based study in Finland. BMJ Open 8, e022752 (2018).

3. Yaffe, K. et al. Sleep-disordered breathing, hypoxia, and risk of mild cognitive impairment and dementia in older women. JAMA 306, 613–9 (2011).

4. Nieto, F. J. et al. Sleep-disordered breathing and cancer mortality: results from the Wisconsin Sleep Cohort Study. Am J Respir Crit Care Med 186, 190–4 (2012).

5. Peppard, P. E. & Hagen, E. W. The Last 25 Years of Obstructive Sleep Apnea Epidemiology-and the Next 25? Am. J. Respir. Crit. Care Med. 197, 310–312 (2018).

6. Redline, S. Genetics of Obstructive Sleep Apnea. in Principles and Practice of Sleep Medicine (ed. Kryger, W. C., Meir H;Roth, Thomas;Dement) 1183–1193 (Saunders, 2011).

7. Kendzerska, T., Gershon, A. S., Hawker, G., Leung, R. S. & Tomlinson, G. Obstructive sleep apnea and risk of cardiovascular events and all-cause mortality: a decade-long historical cohort study. PLoS Med 11, e1001599 (2014).

8. Oldenburg, O. et al. Nocturnal hypoxaemia is associated with increased mortality in stable heart failure patients. Eur Heart J 37, 1695–703 (2016).

9. Gellen, B. et al. Apnea-hypopnea and desaturations in heart failure with reduced ejection fraction: Are we aiming at the right target? Int J Cardiol 203, 1022–8 (2016).

10. Pham, L. V. & Polotsky, V. Y. Genome-Wide Association Studies in Obstructive Sleep Apnea. Will We Catch a Black Cat in a Dark Room? Am J Respir Crit Care Med 194, 789–791 (2016).

11. Cade, B. E. et al. Genetic Associations with Obstructive Sleep Apnea Traits in Hispanic/Latino Americans. Am J Respir Crit Care Med 194, 886–897 (2016).

12. Chen, H. et al. Multiethnic Meta-Analysis Identifies RAI1 as a Possible Obstructive Sleep Apnea-related Quantitative Trait Locus in Men. Am. J. Respir. Cell Mol. Biol. 58, 391–401 (2018).

13. Wang, H. et al. Variants in angiopoietin-2 (ANGPT2) contribute to variation in nocturnal oxyhaemoglobin saturation level. Hum Mol Genet 25, 5244–5253 (2016).

14. Cade, B. E. et al. Associations of Variants In the Hexokinase 1 and Interleukin 18 Receptor Regions with Oxyhemoglobin Saturation During Sleep. PLoS Genet. 15, e1007739 (2019).

15. Wang, H. et al. Admixture mapping identifies novel loci for obstructive sleep apnea in Hispanic/Latino Americans. Hum. Mol. Genet. 28, 675–687 (2019).

16. Gazal, S. et al. Linkage disequilibrium-dependent architecture of human complex traits shows action of negative selection. Nat. Genet. 49, 1421–1427 (2017).

17. Taliun, D. et al. Sequencing of 53,831 diverse genomes from the NHLBI TOPMed Program. Preprint at https://www.biorxiv.org/content/10.1101/563866v1 (2019).

18. Das, S. et al. Next-generation genotype imputation service and methods. Nat. Genet. 48, 1284–1287 (2016).

19. The ARIC Investigators. The Atherosclerosis Risk in Communities (ARIC) Study: design and objectives. The ARIC investigators. Am. J. Epidemiol. 129, 687–702 (1989).

20. Fried, L. P. et al. The Cardiovascular Health Study: design and rationale. Ann Epidemiol 1, 263–76 (1991).

21. Feinleib, M. The Framingham Study: sample selection, follow-up, and methods of analyses. Natl Cancer Inst Monogr 67, 59–64 (1985).

22. Quan, S. F. et al. The Sleep Heart Health Study: design, rationale, and methods. Sleep 20, 1077–85 (1997).

23. Bild, D. E. et al. Multi-Ethnic Study of Atherosclerosis: objectives and design. Am J Epidemiol 156, 871–81 (2002).

24. Chen, X. et al. Racial/Ethnic Differences in Sleep Disturbances: The Multi-Ethnic Study of Atherosclerosis (MESA). Sleep 38, 877–88 (2015).

25. Redline, S. et al. The familial aggregation of obstructive sleep apnea. Am J Respir Crit Care Med 151, 682–7 (1995).

26. Sorlie, P. D. et al. Design and implementation of the Hispanic Community Health Study/Study of Latinos. Ann Epidemiol 20, 629–41 (2010).

27. Redline, S. et al. Sleep-disordered breathing in Hispanic/Latino individuals of diverse backgrounds. The Hispanic Community Health Study/Study of Latinos. Am J Respir Crit Care Med 189, 335–44 (2014).

28. Westbrook, P. R. et al. Description and validation of the apnea risk evaluation system: a novel method to diagnose sleep apnea-hypopnea in the home. Chest 128, 2166–75 (2005).

29. Taylor, H. A. J. et al. Toward resolution of cardiovascular health disparities in African Americans: design and methods of the Jackson Heart Study. Ethn Dis 15, S6–4–17 (2005).

30. Johnson, D. A. et al. Prevalence and correlates of obstructive sleep apnea among African Americans: the Jackson Heart Sleep Study. Sleep 41, (2018).

31. Ng, S. S. S. et al. Validation of Embletta portable diagnostic system for identifying patients with suspected obstructive sleep apnoea syndrome (OSAS). Respirol. Carlton Vic 15, 336–342 (2010).

32. Orwoll, E. et al. Design and baseline characteristics of the osteoporotic fractures in men (MrOS) study–a large observational study of the determinants of fracture in older men. Contemp Clin Trials 26, 569–85 (2005).

33. Blank, J. B. et al. Overview of recruitment for the osteoporotic fractures in men study (MrOS). Contemp Clin Trials 26, 557–68 (2005).

34. Mehra, R. et al. Prevalence and correlates of sleep-disordered breathing in older men: osteoporotic fractures in men sleep study. J Am Geriatr Soc 55, 1356–64 (2007).

35. Hanis, C. L. et al. Diabetes among Mexican Americans in Starr County, Texas. Am J Epidemiol 118, 659–72 (1983).

36. Hanis, C. L. et al. Beyond type 2 diabetes, obesity and hypertension: an axis including sleep apnea, left ventricular hypertrophy, endothelial dysfunction, and aortic stiffness among Mexican Americans in Starr County, Texas. Cardiovasc Diabetol 15, 86 (2016).

37. Choi, J. H. et al. Validation study of portable device for the diagnosis of obstructive sleep apnea according to the new AASM scoring criteria: Watch-PAT 100. Acta Otolaryngol. (Stockh.) 130, 838–843 (2010).

38. Mukherjee, S. et al. Cohort profile: the Western Australian Sleep Health Study. Sleep Breath 16, 205–15 (2012).

39. Redline, S. et al. Methods for obtaining and analyzing unattended polysomnography data for a multicenter study. Sleep Heart Health Research Group. Sleep 21, 759–67 (1998).

40. Whitney, C. W. et al. Reliability of scoring respiratory disturbance indices and sleep staging. Sleep 21, 749–57 (1998).

41. Regier, A. A. et al. Functional equivalence of genome sequencing analysis pipelines enables harmonized variant calling across human genetics projects. Nat. Commun. 9, 4038 (2018).

42. Cingolani, P. et al. A program for annotating and predicting the effects of single nucleotide polymorphisms, SnpEff: SNPs in the genome of Drosophila melanogaster strain w1118; iso-2; iso-3. Fly (Austin) 6, 80–92 (2012).

43. Kang, H. M. et al. Variance component model to account for sample structure in genome-wide association studies. Nat. Genet. 42, 348–354 (2010).

44. Hunt, S. E. et al. Ensembl variation resources. Database J. Biol. Databases Curation 2018, (2018).

45. Dreos, R., Ambrosini, G., Groux, R., Cavin Périer, R. & Bucher, P. The eukaryotic promoter database in its 30th year: focus on non-vertebrate organisms. Nucleic Acids Res. 45, D51–D55 (2017).

46. Yevshin, I., Sharipov, R., Kolmykov, S., Kondrakhin, Y. & Kolpakov, F. GTRD: a database on gene transcription regulation-2019 update. Nucleic Acids Res. (2018). doi: 10.1093/nar/gky1128

47. Rogers, M. F. et al. FATHMM-XF: accurate prediction of pathogenic point mutations via extended features. Bioinforma. Oxf. Engl. 34, 511–513 (2018).

48. di Iulio, J. et al. The human noncoding genome defined by genetic diversity. Nat. Genet. 50, 333–337 (2018).

49. Sundaram, L. et al. Predicting the clinical impact of human mutation with deep neural networks. Nat. Genet. 50, 1161–1170 (2018).

50. Havrilla, J. M., Pedersen, B. S., Layer, R. M. & Quinlan, A. R. A map of constrained coding regions in the human genome. Nat. Genet. 51, 88–95 (2019).

51. Dozmorov, M. G. Epigenomic annotation-based interpretation of genomic data: from enrichment analysis to machine learning. Bioinforma. Oxf. Engl. 33, 3323–3330 (2017).

52. Chèneby, J., Gheorghe, M., Artufel, M., Mathelier, A. & Ballester, B. ReMap 2018: an updated atlas of regulatory regions from an integrative analysis of DNA-binding ChIP-seq experiments. Nucleic Acids Res. 46, D267–D275 (2018).

53. Willer, C. J., Li, Y. & Abecasis, G. R. METAL: fast and efficient meta-analysis of genomewide association scans. Bioinformatics 26, 2190–1 (2010).

54. Barbeira, A. et al. Integrating tissue specific mechanisms into GWAS summary results. (2017). doi:10.1101/045260

55. Zhu, S. et al. GIGSEA: Genotype Imputed Gene Set Enrichment Analysis using GWAS Summary Level Data. Bioinforma. Oxf. Engl. (2018). doi:10.1093/bioinformatics/bty529

56. Cotto, K. C. et al. DGIdb 3.0: a redesign and expansion of the drug-gene interaction database. Nucleic Acids Res. 46, D1068–D1073 (2018).

57. Stelzer, G. et al. The GeneCards Suite: From Gene Data Mining to Disease Genome Sequence Analyses. Curr. Protoc. Bioinforma. 54, 1.30.1–1.30.33 (2016).

58. GTEx Consortium et al. Genetic effects on gene expression across human tissues. Nature 550, 204–213 (2017).

59. Roadmap Epigenomics Consortium et al. Integrative analysis of 111 reference human epigenomes. Nature 518, 317–30 (2015).

60. Martens, J. H. A. & Stunnenberg, H. G. BLUEPRINT: mapping human blood cell epigenomes. Haematologica 98, 1487–9 (2013).

61. Vermunt, M. W. et al. Large-scale identification of coregulated enhancer networks in the adult human brain. Cell Rep 9, 767–79 (2014).

62. Gami, A. S. et al. Obstructive sleep apnea, obesity, and the risk of incident atrial fibrillation. J Am Coll Cardiol 49, 565–71 (2007).

63. Campos-Rodriguez, F. et al. Association between obstructive sleep apnea and cancer incidence in a large multicenter Spanish cohort. Am J Respir Crit Care Med 187, 99–105 (2013).

64. Gozal, D., Ham, S. A. & Mokhlesi, B. Sleep Apnea and Cancer: Analysis of a Nationwide Population Sample. Sleep 39, 1493–500 (2016).

65. Reutrakul, S. & Mokhlesi, B. Obstructive Sleep Apnea and Diabetes: A State of the Art Review. Chest 152, 1070–1086 (2017).

66. Minville, C. et al. Nonalcoholic fatty liver disease, nocturnal hypoxia, and endothelial function in patients with sleep apnea. Chest 145, 525–533 (2014).

67. Corte, T. J. et al. Elevated nocturnal desaturation index predicts mortality in interstitial lung disease. Sarcoidosis Vasc Diffuse Lung Dis 29, 41–50 (2012).

68. Young, T. et al. The occurrence of sleep-disordered breathing among middle-aged adults. N. Engl. J. Med. 328, 1230–1235 (1993).

69. Mohsenin, V. Effects of gender on upper airway collapsibility and severity of obstructive sleep apnea. Sleep Med. 4, 523–529 (2003).

70. Wimms, A., Woehrle, H., Ketheeswaran, S., Ramanan, D. & Armitstead, J. Obstructive Sleep Apnea in Women: Specific Issues and Interventions. BioMed Res. Int. 2016, 1764837 (2016).

71. López-Doménech, G. et al. The Eutherian Armcx genes regulate mitochondrial trafficking in neurons and interact with Miro and Trak2. Nat. Commun. 3, 814 (2012).

72. Serrat, R. et al. The non-canonical Wnt/PKC pathway regulates mitochondrial dynamics through degradation of the arm-like domain-containing protein Alex3. PloS One 8, e67773 (2013).

73. Cartoni, R. et al. The Mammalian-Specific Protein Armcx1 Regulates Mitochondrial Transport during Axon Regeneration. Neuron 92, 1294–1307 (2016).

74. Yoon, J. C. et al. Wnt signaling regulates mitochondrial physiology and insulin sensitivity. Genes Dev. 24, 1507–1518 (2010).

75. Ramezani, R. J. & Stacpoole, P. W. Sleep disorders associated with primary mitochondrial diseases. J. Clin. Sleep Med. JCSM Off. Publ. Am. Acad. Sleep Med. 10, 1233–1239 (2014).

76. Waypa, G. B., Smith, K. A. & Schumacker, P. T. O2 sensing, mitochondria and ROS signaling: The fog is lifting. Mol. Aspects Med. 47–48, 76–89 (2016).

77. Murphy, M. P. How mitochondria produce reactive oxygen species. Biochem. J. 417, 1–13 (2009).

78. Pialoux, V. et al. Effects of exposure to intermittent hypoxia on oxidative stress and acute hypoxic ventilatory response in humans. Am. J. Respir. Crit. Care Med. 180, 1002–1009 (2009).

79. Morgan, B. J., Bates, M. L., Rio, R. D., Wang, Z. & Dopp, J. M. Oxidative stress augments chemoreflex sensitivity in rats exposed to chronic intermittent hypoxia. Respir. Physiol. Neurobiol. 234, 47–59 (2016).

80. MacFarlane, P. M. & Mitchell, G. S. Respiratory long-term facilitation following intermittent hypoxia requires reactive oxygen species formation. Neuroscience 152, 189–197 (2008).

81. Peng, Y.-J., Overholt, J. L., Kline, D., Kumar, G. K. & Prabhakar, N. R. Induction of sensory long-term facilitation in the carotid body by intermittent hypoxia: implications for recurrent apneas. Proc. Natl. Acad. Sci. U. S. A. 100, 10073–10078 (2003).

82. Dan Dunn, J., Alvarez, L. A., Zhang, X. & Soldati, T. Reactive oxygen species and mitochondria: A nexus of cellular homeostasis. Redox Biol. 6, 472–485 (2015).

83. Kim, S. R. et al. NLRP3 inflammasome activation by mitochondrial ROS in bronchial epithelial cells is required for allergic inflammation. Cell Death Dis. 5, e1498 (2014).

84. Kim, S., Lee, Y., Kim, H. & Kim, S. Activation of NLRP3 inflammasome is regulated by mitochondrial ROS via PI3K-HIF-VEGF pathway in acute lung injury. Eur. Respir. J. 46, PA3026 (2015).

85. Moon, J.-S. et al. mTORC1-Induced HK1-Dependent Glycolysis Regulates NLRP3 Inflammasome Activation. Cell Rep 12, 102–15 (2015).

86. Ichimura, H., Parthasarathi, K., Quadri, S., Issekutz, A. C. & Bhattacharya, J. Mechano-oxidative coupling by mitochondria induces proinflammatory responses in lung venular capillaries. J. Clin. Invest. 111, 691–699 (2003).

87. Kim, J. S. et al. Obstructive Sleep Apnea and Subclinical Interstitial Lung Disease in the Multi-Ethnic Study of Atherosclerosis (MESA). Ann. Am. Thorac. Soc. 14, 1786–1795 (2017).

88. Gopisetty, G. & Thangarajan, R. Mammalian mitochondrial ribosomal small subunit (MRPS) genes: A putative role in human disease. Gene 589, 27–35 (2016).

89. Bousquet, P. A. et al. Hypoxia Strongly Affects Mitochondrial Ribosomal Proteins and Translocases, as Shown by Quantitative Proteomics of HeLa Cells. Int. J. Proteomics 2015, 678527 (2015).

90. Seo, M. et al. Genomics and response to long-term oxygen therapy in chronic obstructive pulmonary disease. J. Mol. Med. Berl. Ger. 96, 1375–1385 (2018).

91. Baik, I., Seo, H. S., Yoon, D., Kim, S. H. & Shin, C. Associations of Sleep Apnea, NRG1 Polymorphisms, Alcohol Consumption, and Cerebral White Matter Hyperintensities: Analysis with Genome-Wide Association Data. Sleep 38, 1137–43 (2015).

92. Finigan, J. H. et al. Bronchoalveolar lavage neuregulin-1 is elevated in acute lung injury and correlates with inflammation. Eur. Respir. J. 41, 396–401 (2013).

93. Liu, J., Nethery, D. & Kern, J. A. Neuregulin-1 induces branching morphogenesis in the developing lung through a P13K signal pathway. Exp. Lung Res. 30, 465–478 (2004).

94. Venugopal, R. et al. Inflammasome Inhibition Suppresses Alveolar Cell Permeability Through Retention of Neuregulin-1 (NRG-1). Cell. Physiol. Biochem. Int. J. Exp. Cell. Physiol. Biochem. Pharmacol. 36, 2012–2024 (2015).

95. Lederer, D. J., Jelic, S., Basner, R. C., Ishizaka, A. & Bhattacharya, J. Circulating KL-6, a biomarker of lung injury, in obstructive sleep apnoea. Eur Respir J 33, 793–6 (2009).

96. Paatero, I. et al. Interaction with ErbB4 promotes hypoxia-inducible factor-1α signaling. J. Biol. Chem. 287, 9659–9671 (2012).

97. Kyotani, Y. et al. Intermittent hypoxia induces the proliferation of rat vascular smooth muscle cell with the increases in epidermal growth factor family and erbB2 receptor. Exp. Cell Res. 319, 3042–3050 (2013).

98. Gamazon, E. R., Konkashbaev, A., Derks, E. M., Cox, N. J. & Lee, Y. Evidence of selection on splicing-associated loci in human populations and relevance to disease loci mapping. Sci. Rep. 7, 5980 (2017).

99. Stauffer, T. P., Guerini, D. & Carafoli, E. Tissue distribution of the four gene products of the plasma membrane Ca2+ pump. A study using specific antibodies. J Biol Chem 270, 12184–90 (1995).

100. Schuh, K. et al. Regulation of vascular tone in animals overexpressing the sarcolemmal calcium pump. J Biol Chem 278, 41246–52 (2003).

101. Kell, A. M. & Gale, M. RIG-I in RNA virus recognition. Virology 479–480, 110–121 (2015).

102. Gharib, S. A., Hayes, A. L., Rosen, M. J. & Patel, S. R. A pathway-based analysis on the effects of obstructive sleep apnea in modulating visceral fat transcriptome. Sleep 36, 23–30 (2013).

103. Herriges, J. C. et al. FGF-Regulated ETV Transcription Factors Control FGF-SHH Feedback Loop in Lung Branching. Dev. Cell 35, 322–332 (2015).

104. Wollenick, K. et al. Synthetic transactivation screening reveals ETV4 as broad coactivator of hypoxia-inducible factor signaling. Nucleic Acids Res. 40, 1928–1943 (2012).

105. Shu, W., Yang, H., Zhang, L., Lu, M. M. & Morrisey, E. E. Characterization of a new subfamily of winged-helix/forkhead (Fox) genes that are expressed in the lung and act as transcriptional repressors. J. Biol. Chem. 276, 27488–27497 (2001).

106. Chokas, A. L. et al. Foxp1/2/4-NuRD interactions regulate gene expression and epithelial injury response in the lung via regulation of interleukin-6. J. Biol. Chem. 285, 13304–13313 (2010).

107. Xu, S. et al. Foxp2 regulates anatomical features that may be relevant for vocal behaviors and bipedal locomotion. Proc. Natl. Acad. Sci. U. S. A. 115, 8799–8804 (2018).

108. Stanić, D., Dhingra, R. R. & Dutschmann, M. Expression of the transcription factor FOXP2 in brainstem respiratory circuits of adult rat is restricted to upper-airway pre-motor areas. Respir. Physiol. Neurobiol. 250, 14–18 (2018).

109. Burgold, T. et al. The H3K27 demethylase JMJD3 is required for maintenance of the embryonic respiratory neuronal network, neonatal breathing, and survival. Cell Rep. 2, 1244–1258 (2012).

110. Smith, J. C., Ellenberger, H. H., Ballanyi, K., Richter, D. W. & Feldman, J. L. Pre-Bötzinger complex: a brainstem region that may generate respiratory rhythm in mammals. Science 254, 726–729 (1991).

111. Chen, Y. et al. Global Histone H3K23/H3K36 Hypoacetylation and HDAC1 Up-regulation Are Associated with Disease Severity and Adverse Consequences in Obstructive Sleep Apnea Patients. Am J Respir Crit Care Med 197, A6411 (2018).

112. Satoh, T. et al. The Jmjd3-Irf4 axis regulates M2 macrophage polarization and host responses against helminth infection. Nat. Immunol. 11, 936–944 (2010).

113. De Santa, F. et al. The histone H3 lysine-27 demethylase Jmjd3 links inflammation to inhibition of polycomb-mediated gene silencing. Cell 130, 1083–1094 (2007).

114. Li, Q. et al. Stage-dependent and locus-specific role of histone demethylase Jumonji D3 (JMJD3) in the embryonic stages of lung development. PLoS Genet. 10, e1004524 (2014).

115. Shalaby, N. A. et al. JmjC domain proteins modulate circadian behaviors and sleep in Drosophila. Sci. Rep. 8, 815 (2018).

116. Shmakova, A., Batie, M., Druker, J. & Rocha, S. Chromatin and oxygen sensing in the context of JmjC histone demethylases. Biochem. J. 462, 385–395 (2014).

117. Hancock, R. L., Masson, N., Dunne, K., Flashman, E. & Kawamura, A. The Activity of JmjC Histone Lysine Demethylase KDM4A is Highly Sensitive to Oxygen Concentrations. ACS Chem. Biol. 12, 1011–1019 (2017).

118. Yan, S. F. et al. Tissue factor transcription driven by Egr-1 is a critical mechanism of murine pulmonary fibrin deposition in hypoxia. Proc. Natl. Acad. Sci. U. S. A. 95, 8298–8303 (1998).

119. Thomason, H. A., Dixon, M. J. & Dixon, J. Facial clefting in Tp63 deficient mice results from altered Bmp4, Fgf8 and Shh signaling. Dev. Biol. 321, 273–282 (2008).

120. Robison, J. G. & Otteson, T. D. Increased prevalence of obstructive sleep apnea in patients with cleft palate. Arch. Otolaryngol. Head Neck Surg. 137, 269–274 (2011).

121. Obeidat, M. et al. GSTCD and INTS12 regulation and expression in the human lung. PloS One 8, e74630 (2013).

