## Supplemental Figures for "Whole-Genome Association Analyses of Sleep-disordered Breathing Phenotypes in the NHLBI TOPMed Program"

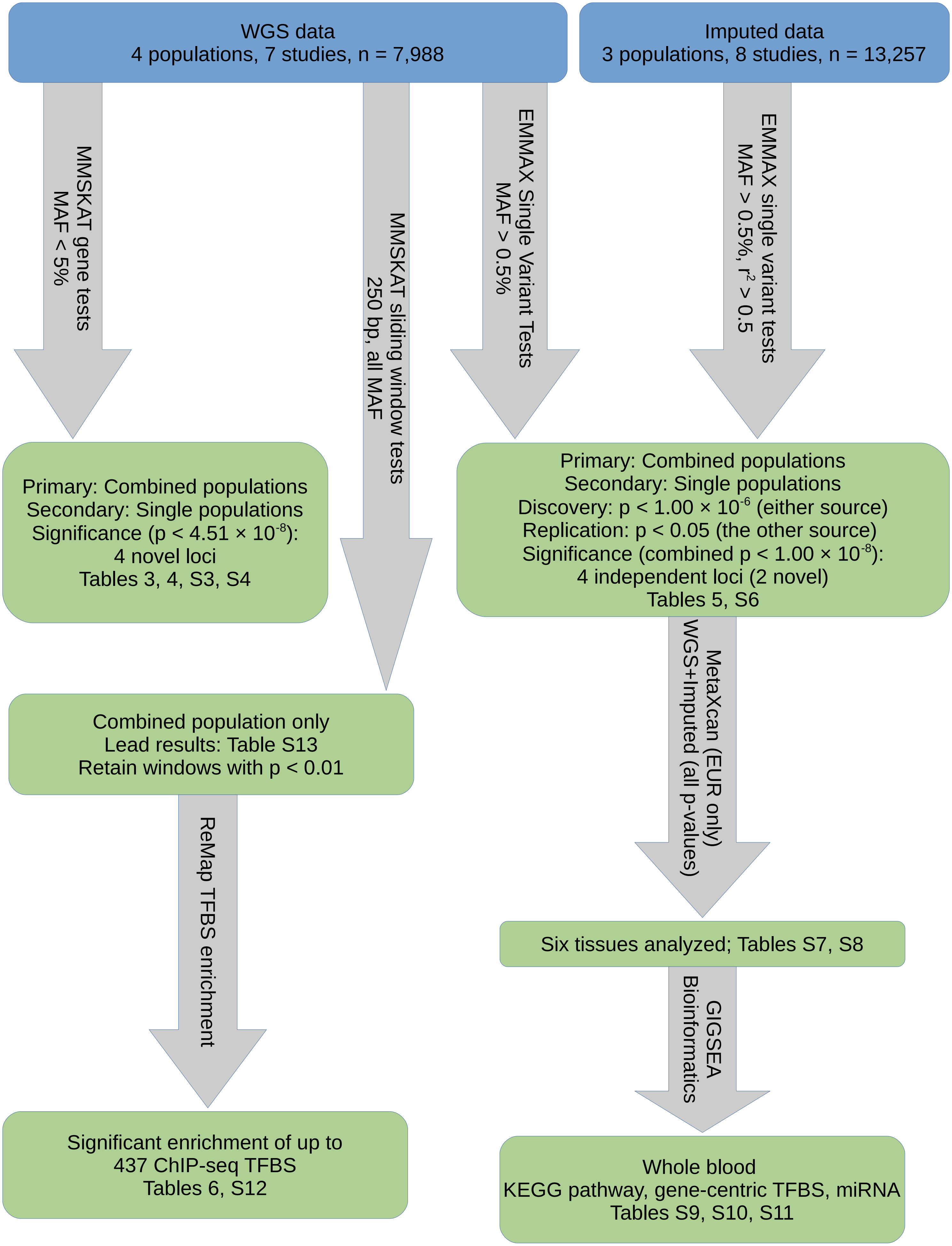
WGS data
4 populations, 7 studies, n = 7,988
Imputed data
3 populations, 8 studies, n = 13,257
MMSKAT gene tests
MAF < 5%
MMSKAT sliding window tests
250 bp, all MAF
EMMAX Single Variant Tests
MAF > 0.5%
EMMAX single variant tests
MAF > 0.5%, r2 > 0.5
Primary: Combined populations
Secondary: Single populations
Significance (p < 4.51 × 10-8):
4 novel loci
Tables 3, 4, S3, S4
Primary: Combined populations
Secondary: Single populations
Discovery: p < 1.00 × 10-6 (either source)
Replication: p < 0.05 (the other source)
Significance (combined p < 1.00 × 10-8):
4 independent loci (2 novel)
Tables 5, S6
MetaXcan (EUR only)
WGS+Imputed (all p-values)
Combined population only
Lead results: Table S13
Retain windows with p < 0.01
Six tissues analyzed; Tables S7, S8
ReMap TFBS enrichment
GIGSEA
Bioinformatics
Significant enrichment of up to
437 ChIP-seq TFBS
Tables 6, S12
Whole blood
KEGG pathway, gene-centric TFBS, miRNA
Tables S9, S10, S11

### Supplementary Figure 2 NRG1 Locus Models Manhattan and QQ Plots

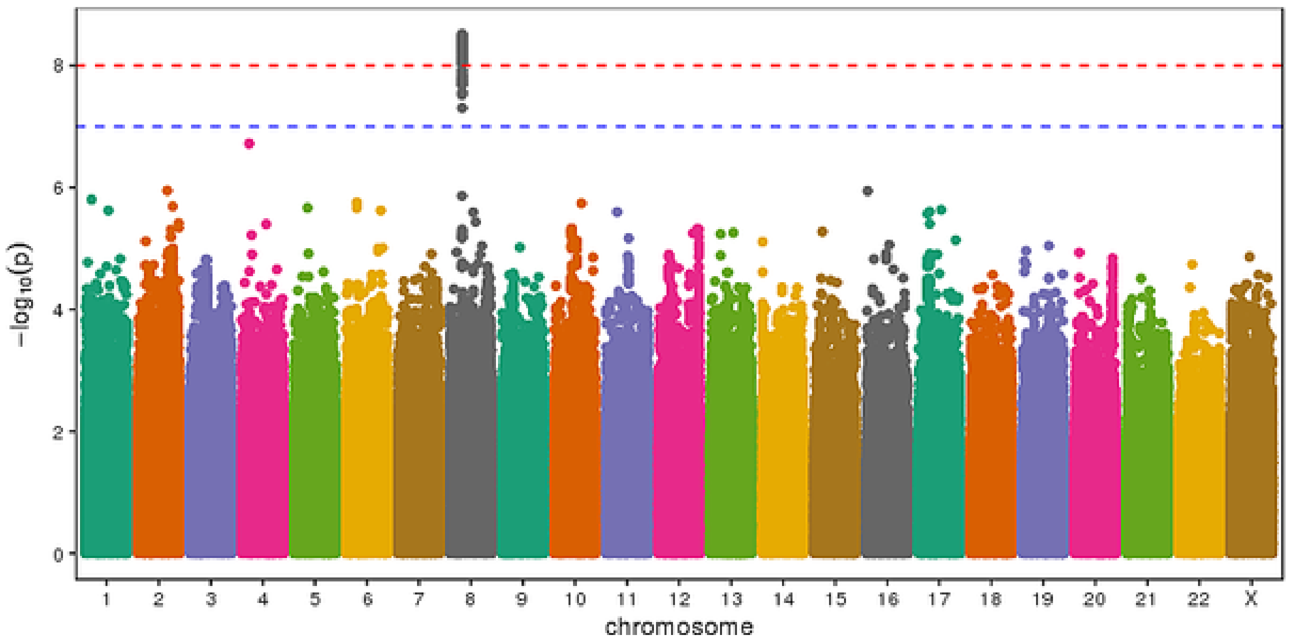


AHI European-American/Australians

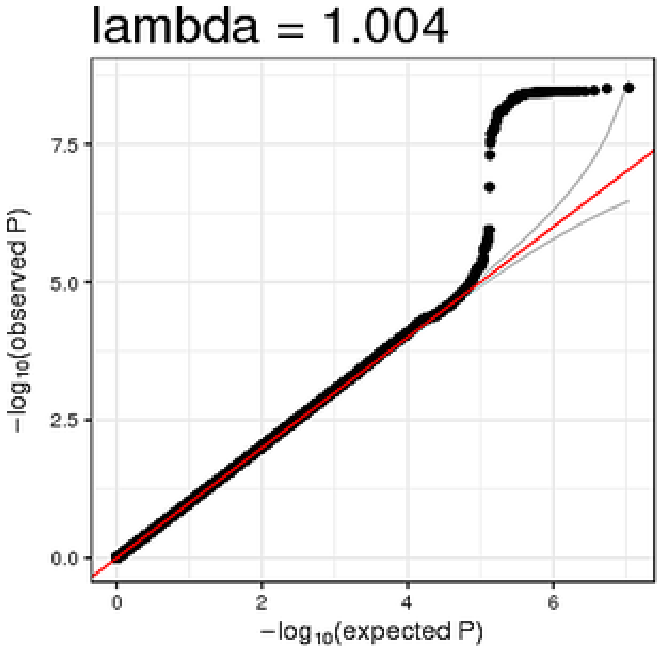


AHI European-American/Australians

### Supplementary Figure 3 SLC45A2 Locus Models Manhattan and QQ Plots

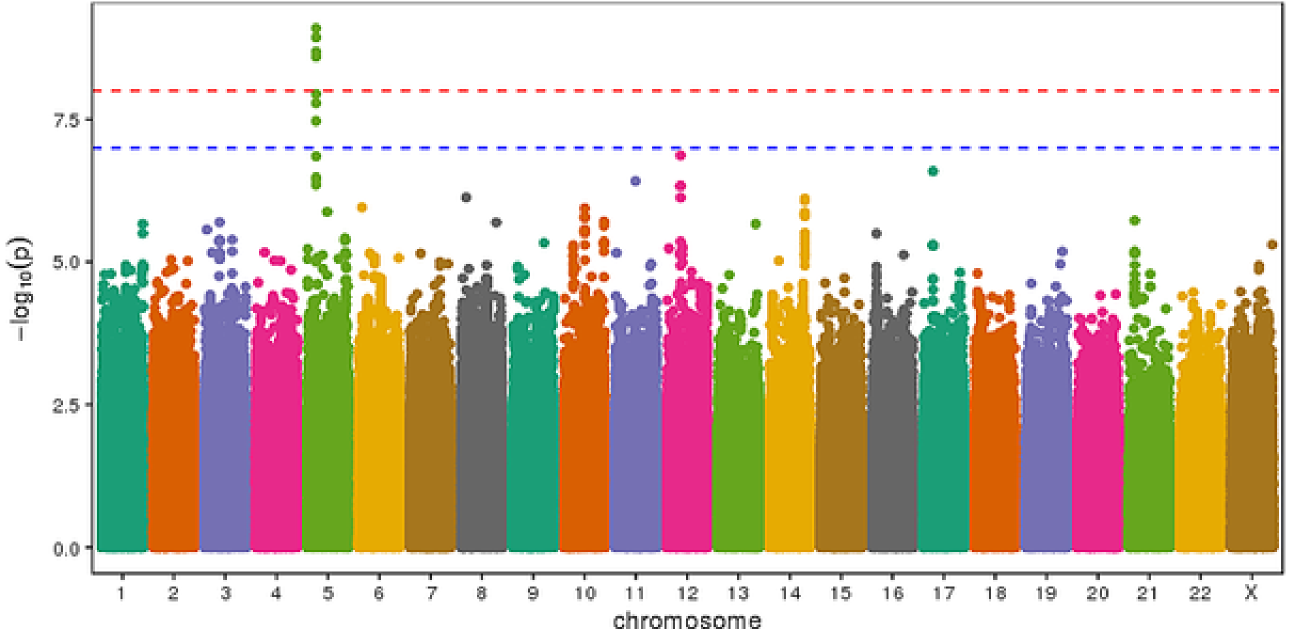


Average SpO2 European-American/Australians

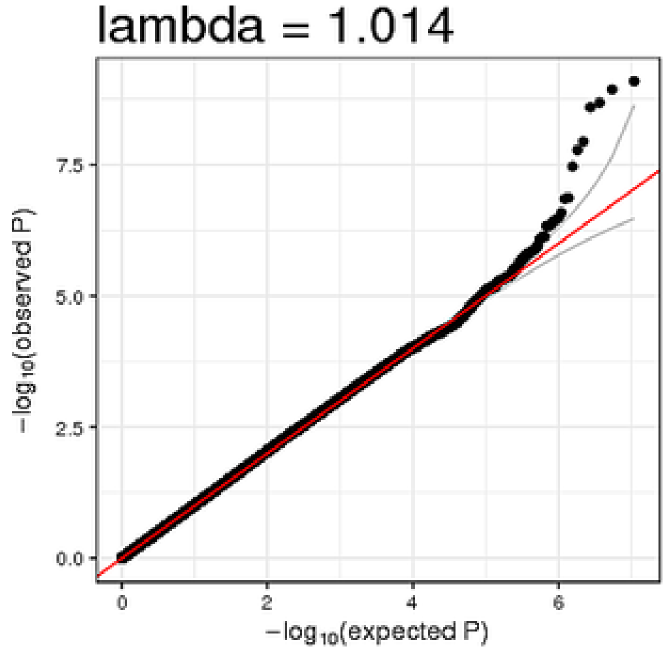


Average SpO2 European-American/Australians

### Supplementary Figure 4 IL18RAP Locus Models Manhattan and QQ Plots

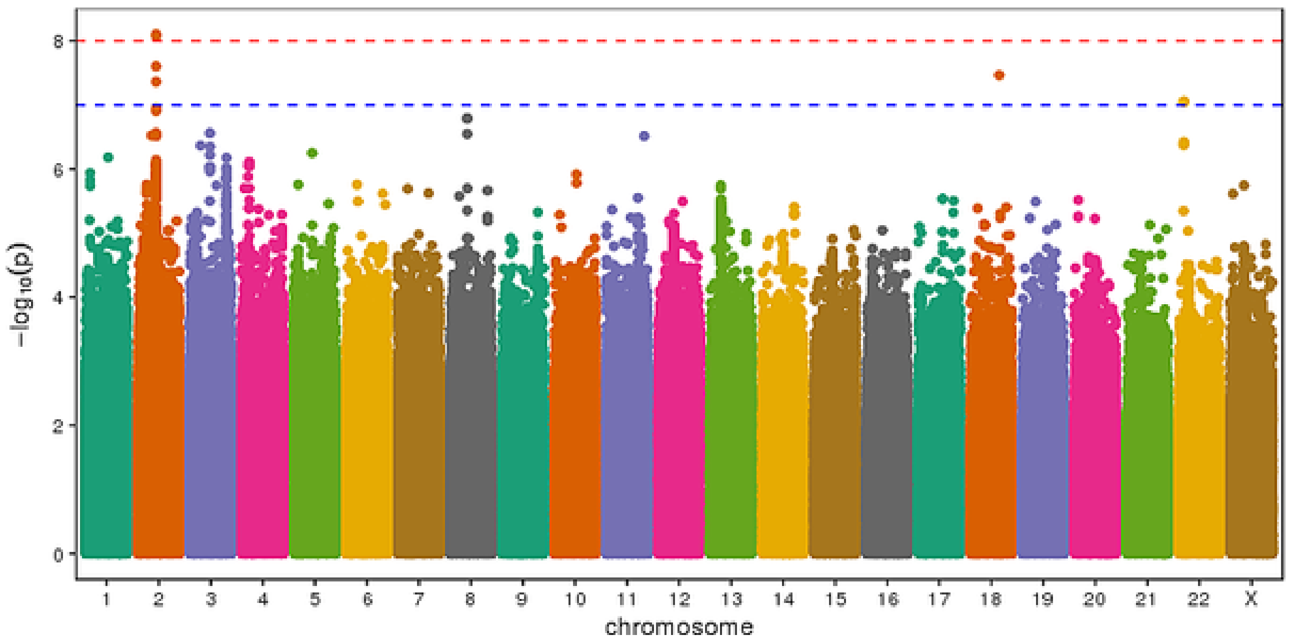


Minimum SpO2 Combined

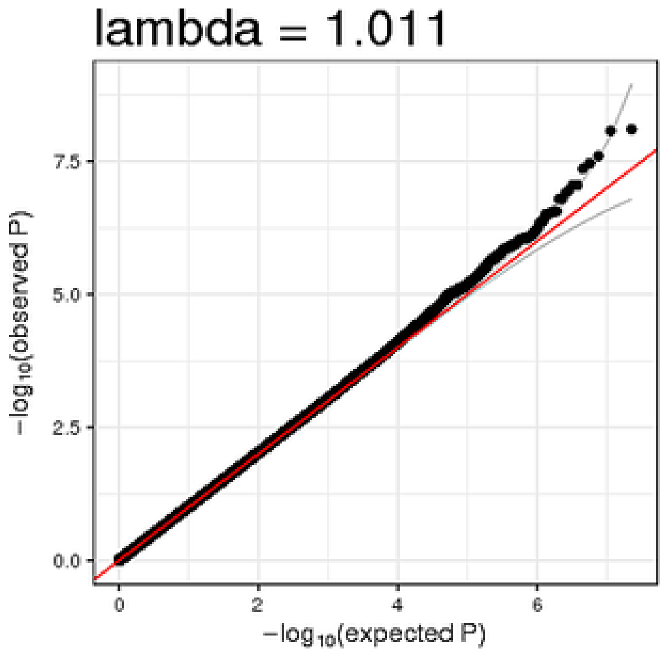


Minimum SpO2 Combined

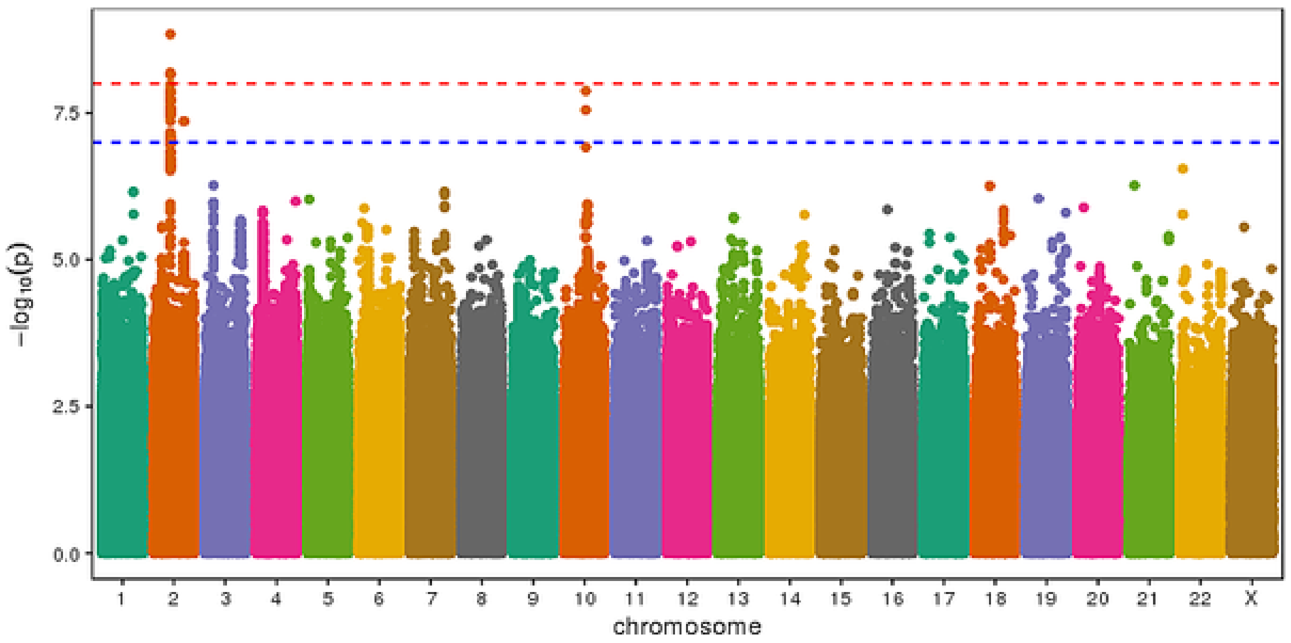


Minimum SpO2 Males

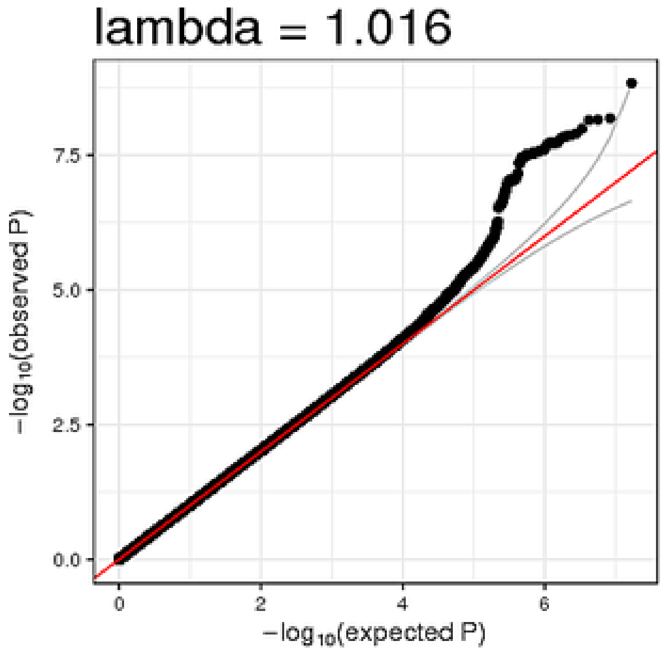


Minimum SpO2 Males

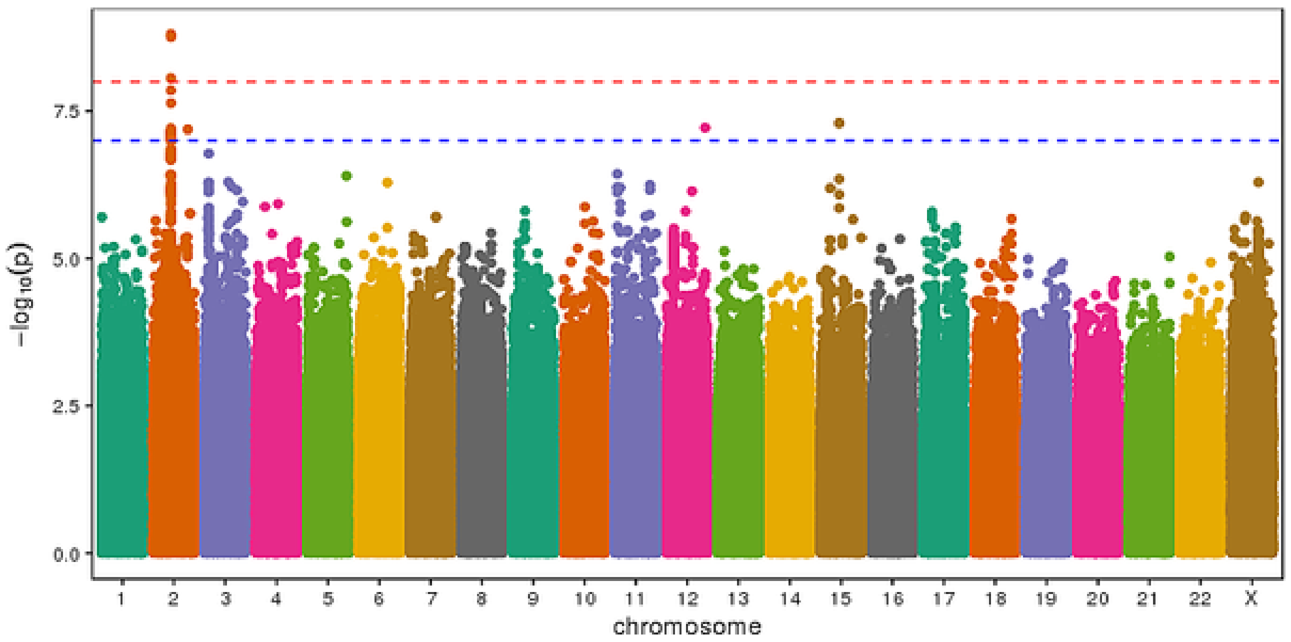


Average Desaturation Combined

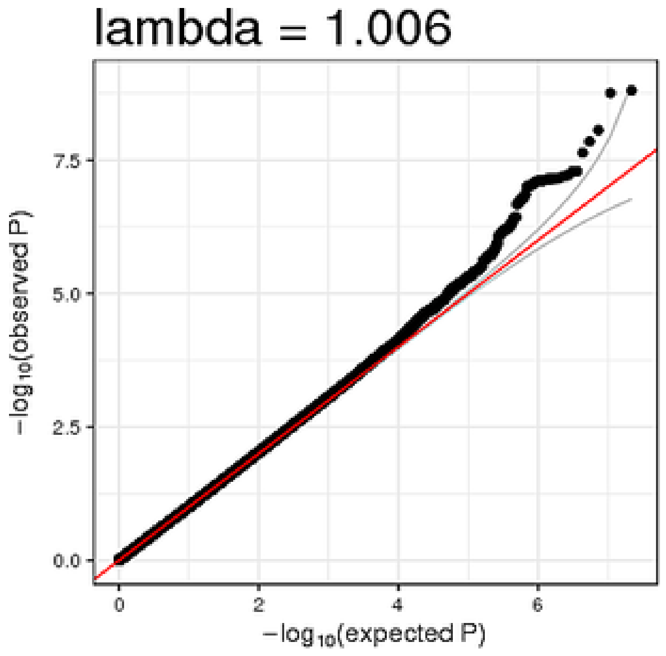


Average Desaturation Combined

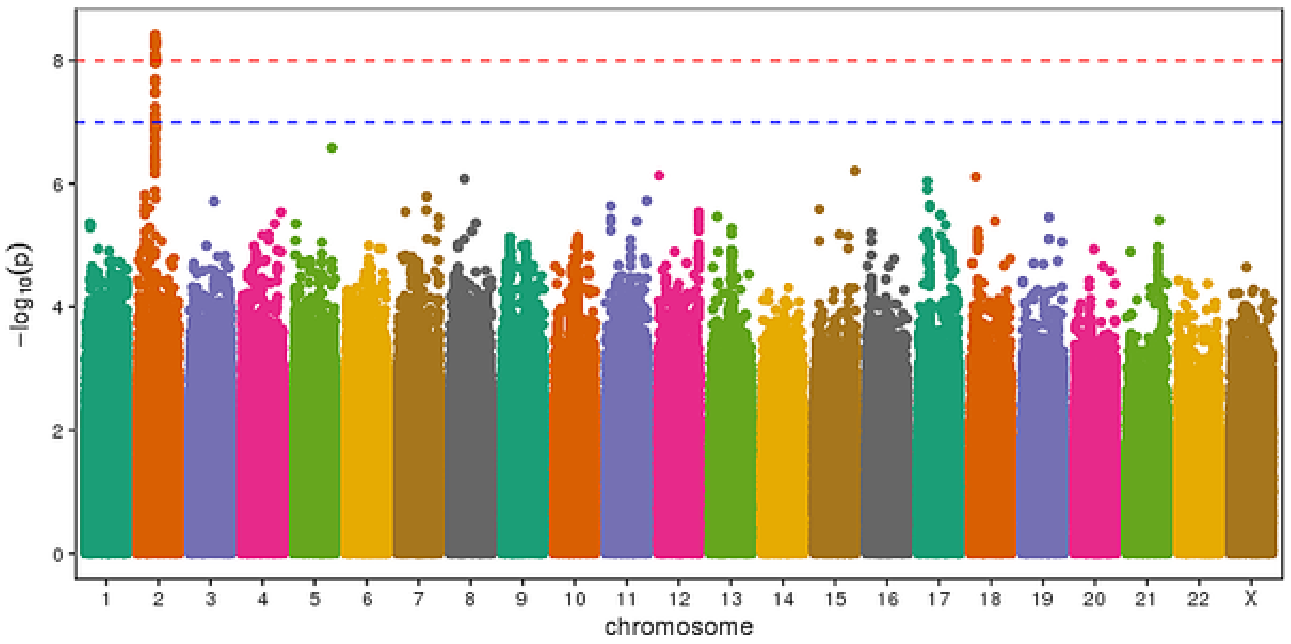


Average Desaturation Males

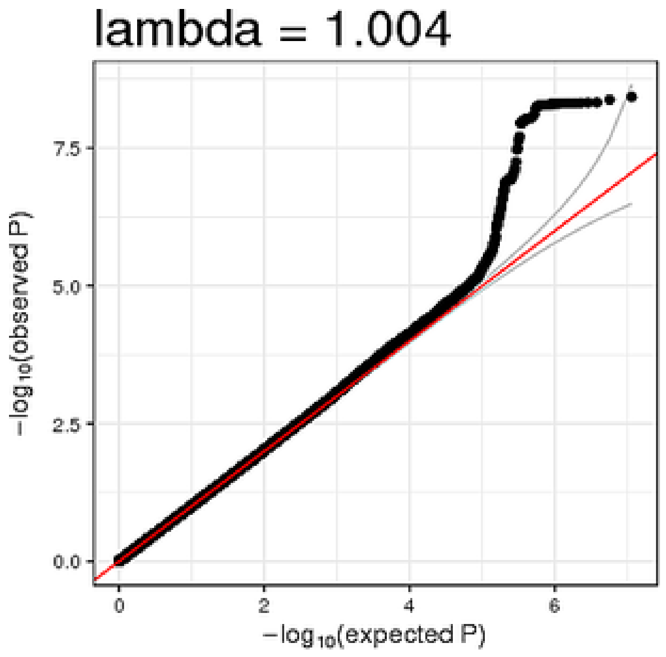


Average Desaturation Males

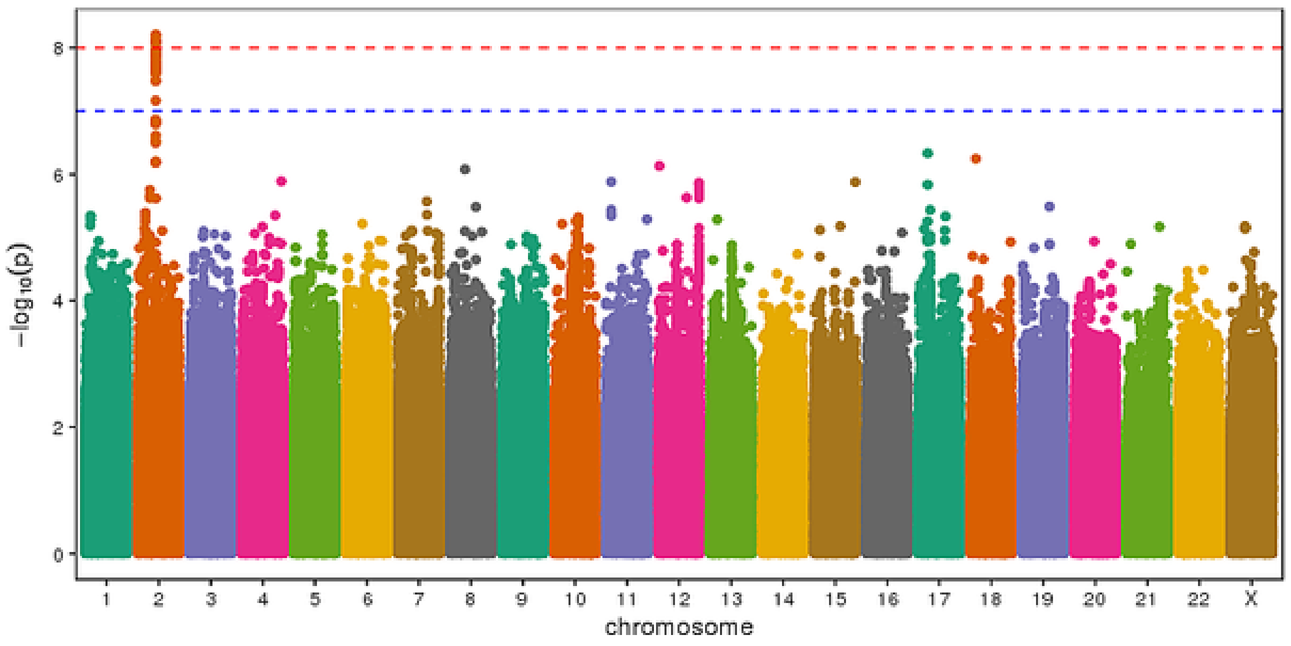


Average Desaturation European-American/Australian Males

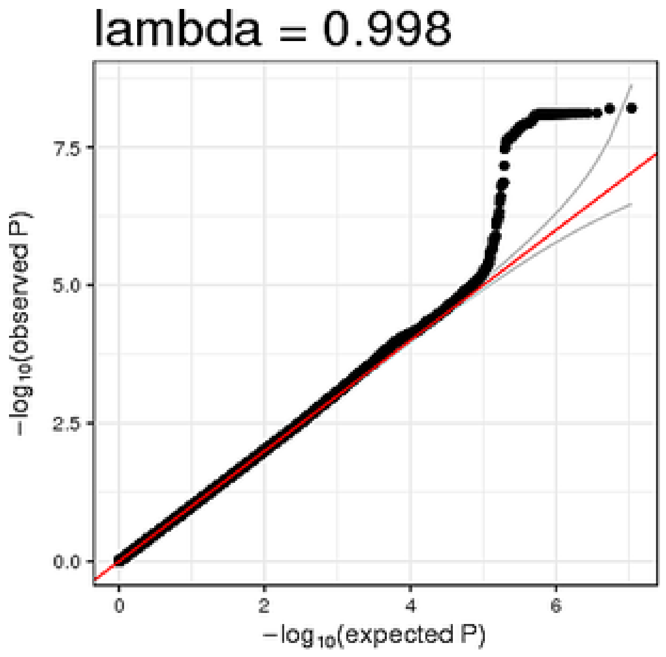


Average Desaturation European-American/Australian Males

### Supplementary Figure 5 ATP2B4 Locus Models Manhattan and QQ Plots

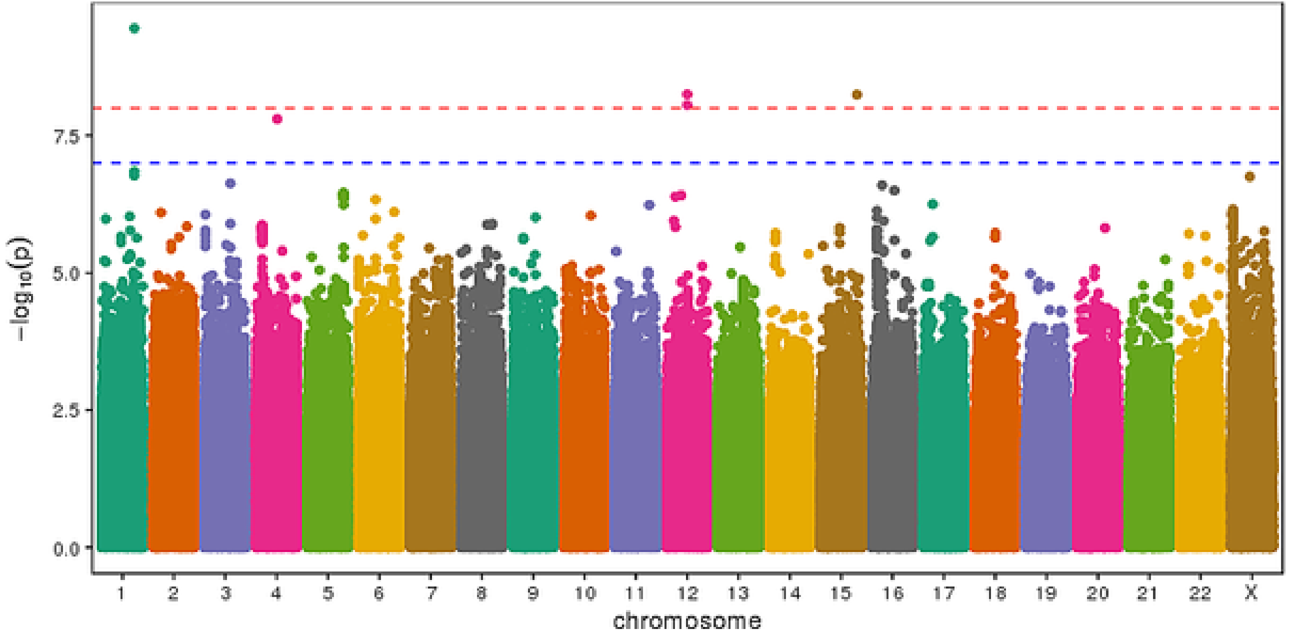


Per90 Hispanic/Latino Americans

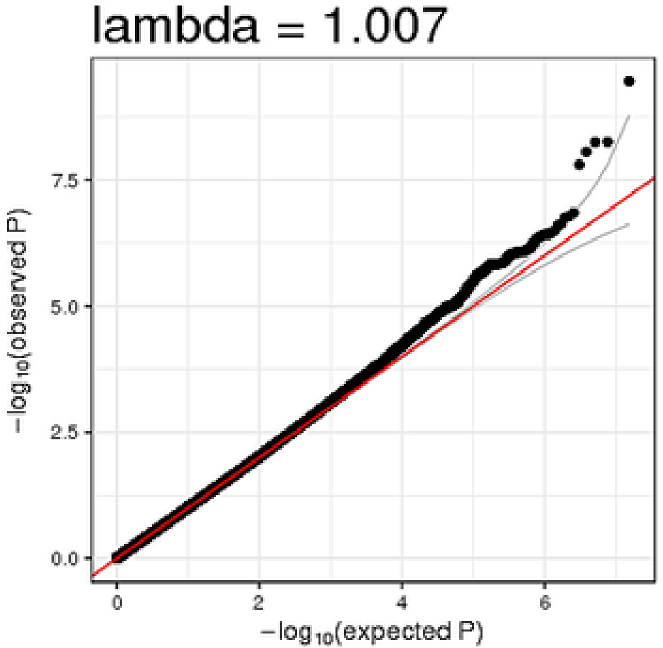


Per90 Hispanic/Latino Americans
